# Design of efficient artificial enzymes using crystallographically-enhanced conformational sampling

**DOI:** 10.1101/2023.11.01.564846

**Authors:** Rojo V. Rakotoharisoa, Behnoush Seifinoferest, Niayesh Zarifi, Jack D.M. Miller, Joshua M. Rodriguez, Michael C. Thompson, Roberto A. Chica

## Abstract

The ability to create efficient artificial enzymes for any chemical reaction is of great interest. Here, we describe a computational design method for increasing catalytic efficiency of *de novo* enzymes to a level comparable to their natural counterparts without relying on directed evolution. Using structural ensembles generated from dynamics-based refinement against X-ray diffraction data collected from crystals of Kemp eliminases HG3 (*k*_cat_/*K*_M_ 125 M^−1^ s^−1^) and KE70 (*k*_cat_/*K*_M_ 57 M^−1^ s^−1^), we design from each enzyme ≤10 sequences predicted to catalyze this reaction more efficiently. The most active designs display *k*_cat_/*K*_M_ values improved by 100–250-fold, comparable to mutants obtained after screening thousands of variants in multiple rounds of directed evolution. Crystal structures show excellent agreement with computational models. Our work shows how computational design can generate efficient artificial enzymes by exploiting the true conformational ensemble to more effectively stabilize the transition state.

## Main text

The ability to create efficient artificial enzymes for any desired chemical reaction is of great interest as it would open the door to a broad range of applications in industry and medicine. Algorithms for *de novo* enzyme design^1,2^ can be used to create artificial enzymes for reactions for which no naturally occurring enzyme is known. While this computational design process has been used to produce *de novo* enzymes for various model organic transformations^3–7^, downstream directed evolution has been necessary to convert these enzyme prototypes into catalysts as efficient as their natural counterparts^8–13^. However, directed evolution is costly, time-consuming and impractical for many enzymatic reactions of interest for which high-throughput screening assays are unavailable. Therefore, a computational design procedure to enhance catalytic efficiency of *de novo* enzymes would be beneficial as it could bypass directed evolution and accelerate the creation of efficient artificial enzymes.

Structural analyses of high activity variants evolved from *de novo* enzymes revealed several mechanisms by which catalytic function can be augmented. These mechanisms include optimization of hydrogen-bond geometries between catalytic residues and the transition state^8^, enhancement of active-site complementarity to the transition state^8,9,14^, improvement of active-site preorganization^14^, and enrichment of catalytically competent substates in the conformational ensemble^15–17^. These structural effects are challenging to design using current enzyme design protocols^3,7,9^ because they are not sufficiently precise^8^ and do not consider the dynamic nature of enzyme structures^15^. Recently, new methodologies have been developed that take advantage of the spatiotemporal averaging inherent to X-ray crystallography – crystallographic data represent billions of crystallized molecules that are all sampling different conformations during data collection – to model the conformational ensembles of proteins with high fidelity^18,19^. New design strategies that could exploit this experimental information should facilitate the design of efficient artificial enzymes by enabling accurate modeling of catalytically productive conformational substates.

Here, we report a computational procedure for enhancing the catalytic efficiency of *de novo* enzymes that exploits crystallographically-derived ensemble models for increased accuracy. Starting from the non-cryogenic crystal structure of *de novo* Kemp eliminase HG3^7,14^, we generate a structural ensemble fitted to the diffraction data^18,20^ and use it to design mutant sequences predicted to efficiently catalyze the Kemp elimination. We filter these sequences using a variety of structural criteria aiming to reproduce molecular mechanisms known to augment enzyme catalysis, and experimentally characterize 10 sequences. The most active design displays a catalytic efficiency improved by approximately 250-fold, a result comparable to those obtained by seven rounds of directed evolution and screening of >7000 variants^8^. We solve crystal structures of several high-activity variants, which show excellent agreement with the computational models and confirm that catalytic efficiency is enhanced via the intended molecular mechanisms. We also demonstrate the general applicability of our approach by applying it to an unrelated *de novo* Kemp eliminase, KE70^3^, which yields comparable results. Our work shows how ensemble-based design can generate efficient artificial enzymes by exploiting information derived from X-ray crystallography to identify productive substates for efficient catalysis.

## Results

### Ensemble-based enzyme design

To create efficient *de novo* enzymes without relying on iterative rounds of mutagenesis and high-throughput screening, we implemented an ensemble-based computational enzyme design pipeline that uses protein backbones generated by ensemble refinement of diffraction data^18^ as templates for design (Figure 1). Because the diffraction data and calculated electron density maps are derived from crystals that contain billions of molecules, which are all sampling different conformational states throughout data collection, the resulting ensemble models are expected to represent the entire equilibrium conformational ensemble of the target protein fold^21^. Thus, we postulated that performing crystallographic ensemble refinement on a low activity *de novo* enzyme would allow sampling of conformational substates that can more accurately accommodate the transition state and its interactions with catalytic residues than the single conformation modeled using traditional refinement methods, thereby facilitating the design of improved active sites. To test our hypothesis, we applied ensemble-based design to increase catalytic efficiency of the *de novo* enzyme HG3^7^ that catalyzes the Kemp elimination (Supplementary Figure 1a), a well-established model organic transformation for benchmarking computational enzyme design methods. We selected HG3 as a case study for three reasons. First, X-ray diffraction data collected at non-cryogenic temperatures (277 K) were available^14^, which is ideal for ensemble refinement because cryocooling is known to narrow the conformational ensemble of crystallized proteins^22^, reducing the number of conformations that are represented in the ensemble-averaged data. Second, these data were collected from crystals with a bound transition-state analogue that is resolved in the electron density^14^, allowing inclusion of the transition state in the ensemble models. Finally, HG3 was subjected to an extensive directed evolution campaign that increased catalytic efficiency by almost three orders of magnitude^8^, allowing us to compare results with those from purely experimental approaches.

**Figure 1.**
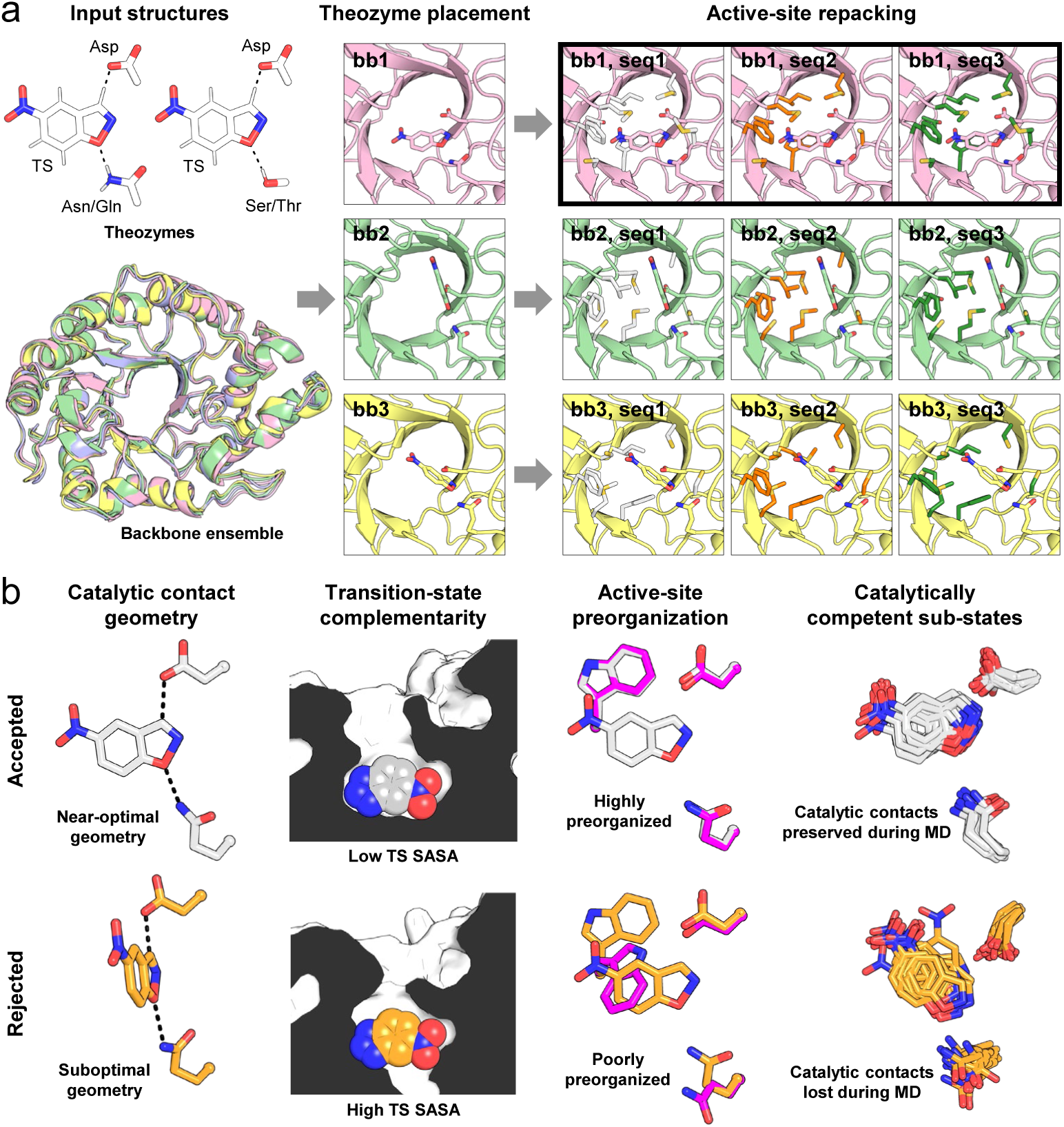
Ensemble-based computational enzyme design. (a) This procedure requires as inputs one or more theozymes, which are combinations of catalytic groups arranged in a unique geometry to stabilize a transition state (TS), and a backbone ensemble that approximates the intrinsic flexibility of the target protein fold. The algorithm proceeds in two stages: placement of each theozyme onto every member of the backbone ensemble (bb1‒3), followed by repacking of active-site residues to identify sequences (seq1‒3) that can further stabilize the TS. During repacking, both the identity and conformation of designed active-site residues are allowed to vary, and the TS is rotated and translated to reduce steric clashes with repacked side chains. In this toy example, only the theozyme containing a Gln hydrogen-bond donor is shown for clarity. After active-site repacking, designs from a single theozyme/template combinations are selected based on energy (bold outline) and filtered using structural criteria. (b) Filtering of designs is performed to reproduce mechanisms known to contribute to efficient enzyme catalysis. Accepted designs are those that have (1) near-ideal contact geometries between the TS and catalytic side chains (dashed lines), (2) active sites with high structural complementarity to the TS, as calculated by low TS solvent-accessible surface area (SASA), (3) highly preorganized active sites with most designed residues predicted to adopt the same rotameric configuration in the presence (white/orange) and absence (magenta) of TS, and (4) enriched catalytically competent substates, which are snapshots that preserve catalytic contacts over the course of short molecular dynamics (MD) simulations.

We generated an ensemble of 84 backbone templates (Supplementary Table 1) from the crystal structure of HG3 bound with the 6-nitrobenzotriazole (6NT, Supplementary Figure 1b) transition-state analogue (PDB ID: 5RGA^14^) and used these structures for placement of four theozymes containing an Asp residue as catalytic base, and either a Gln, Asn, Ser or Thr to act as hydrogen-bond donor for stabilizing negative charge buildup on the phenolic oxygen at the transition state (Figure 1a). We kept the base position fixed to that of HG3 (D127) but allowed the algorithm to place the hydrogen-bond donor at multiple positions in the active site given that no amino acid residue in HG3 forms this catalytic contact with 6NT^14^. We then repacked the active site around the most stable theozyme placed on each backbone template by optimizing the identity and side-chain conformation of neighbouring residues to maximize beneficial packing interactions with the transition state, which has been shown to contribute to increased catalytic activity^8^.

After repacking, we filtered designs using a series of structural metrics aiming to reproduce mechanisms known to augment enzyme catalysis (Figure 1b, Supplementary Table 2). First, we measured angles defining the hydrogen bond between D127 and the transition state for the top-scoring design obtained on each backbone template, and selected sequences from the single backbone template that gave the most stable design with angles that were closest to the optimal geometries calculated for acetamide dimers^23^ (Supplementary Figure 2, Supplementary Table 3). The selected backbone, which we name bb05, yielded sequences that contain the Q50 hydrogen-bond donor that emerged during evolution of HG3 and that is present in the highest activity variants of this enzyme family^8^. Second, we evaluated active-site complementarity to the transition state by calculating the solvent-accessible surface area of this ligand in the computational models generated for 1000 sequences designed on backbone bb05. Only designs where ≥95% of the transition state surface is buried were kept. Third, we evaluated active-site preorganization by calculating the percentage of designed active-site residues that were predicted to adopt similar rotamers in the absence and presence of the transition state, which we consider to be preorganized for catalysis. Sequences whose active-sites were predicted to be >70% preorganized were kept. We also eliminated sequences with calculated energies over 10 kcal mol^−1^ higher than that of an HG3 single point mutant containing the D127/Q50 catalytic dyad, as well as those that contained more than four methionines or more than one cysteine in the active site to avoid excessive conformational heterogeneity and prevent formation of disulfide bonds, respectively. This filtering process narrowed the original list of 1000 designs obtained from backbone bb05 down to 55 designs. A final filtering step consisting of a 20-ns molecular dynamics simulation was used to identify designs that were enriched in catalytically-competent conformational substates, which we define as snapshots that make the designed catalytic contacts (Supplementary Figure 3). This final filtering resulted in nine sequences containing four or five mutations each (Table 1), which we experimentally characterized.

**Table 1.** Kinetic parameters of Kemp eliminases.

| Enzyme | Mutations <sup>a</sup> | #<br>Mutations <sup>a</sup> | $k_{cat}$<br>(s<sup>-1</sup>) <sup>b</sup> | $K_M$<br>(mM) <sup>b</sup> | $k_{cat}/K_M$<br>(M<sup>-1</sup> s<sup>-1</sup>) <sup>b</sup> | Relative<br>$k_{cat}/K_M$ |
| --- | --- | --- | --- | --- | --- | --- |
| **HG series – Controls** |  |  |  |  |  |  |
| **HG3** | – | – | N.D. | N.D. | 125 ± 8<br>(146 ± 6) <sup>c</sup> | 1 |
| **HG3.3b** | V6I K50H M84C S89R<br>Q90D A125N | 6 | N.D. | N.D. | 3400 ± 700<br>(2200 ± 100) <sup>c</sup> | 27 |
| **HG3-K50Q/M84C** | K50Q M84C | 2 | N.D. | N.D. | 4300 ± 300 | 34 |
| **HG3.7** | V6I Q37K K50Q M84C<br>S89R Q90H A125N | 7 | N.D. | N.D. | 24000 ± 3000<br>(27000 ± 2000) <sup>c</sup> | 192 |
| **HG series – Designs** |  |  |  |  |  |  |
| **HG378** | A21C K50Q Q90L A125V<br>W275V | 5 | N.D. | N.D. | 70 ± 5 | 0.6 |
| **HG57** | A21C K50Q Q90L A125V | 4 | N.D. | N.D. | 120 ± 8 | 1 |
| **HG44** | A21C K50Q Q90F A125V | 4 | N.D. | N.D. | 250 ± 10 | 2 |
| **HG603** | K50Q Q90F A125V W275V | 4 | N.D. | N.D. | 800 ± 10 | 6.4 |
| **HG493** | K50Q Q90F A125T W275T | 4 | N.D. | N.D. | 2100 ± 100 | 17 |
| **HG185** | A21C K50Q Q90F A125T<br>W275V | 5 | N.D. | N.D. | 2300 ± 100 | 18 |
| **HG198** | K50Q M84C A125T F267M | 4 | N.D. | N.D. | 8000 ± 300 | 64 |
| **HG259** | K50Q M84C Q90L A125T<br>F267M | 5 | N.D. | N.D. | 13000 ± 400 | 104 |
| **HG630** | K50Q M84C A125T F267M<br>W275V | 5 | N.D. | N.D. | 15000 ± 500 | 120 |
| **Without MD filtering <sup>d</sup>** |  |  |  |  |  |  |
| **HG649** | K50Q M84C Q90F A125T<br>F267M W275V | 6 | N.D. | N.D. | 32000 ± 3000 | 256 |
| **KE70 series – Controls** |  |  |  |  |  |  |
| **KE70** | – | – | N.D. | N.D. | 57 ± 3<br>(78 ± 14) <sup>c</sup> | 1 |
| **KE70-S20a/A240a <sup>e</sup>** | S20a A240a | 2 | N.D. | N.D. | 150 ± 7 | 2.6 |
| **R2 7/12F** | D23G, Y48F, D212E,<br>H251Y | 4 | 0.50 ± 0.01 | 1.00 ± 0.06 | 500 ± 30<br>(1330 ± 13) <sup>c</sup> | 9 |
| **R4 8/5A <sup>e</sup>** | S20a, K29N, Y48F, W72C,<br>H166Y, K197N, A204V,<br>G240a | 8 | 3.0 ± 0.1 | 1.0 ± 0.1 | 3000 ± 300<br>(4380 ± 350) <sup>c</sup> | 53 |
| **KE70 series – Designs <sup>f</sup>** |  |  |  |  |  |  |
| **KE701 <sup>e</sup>** | H16D S20a D45Q Y48F<br>W72C G101S S138Q H166N<br>V168A A204V A240a | 11 | 0.040 ± 0.001 | 0.10 ± 0.02 | 400 ± 80 | 7 |
| **KE703 <sup>e</sup>** | H16D S20a D45N Y48F<br>W72V G101M S138Q<br>H166N V168A A204V<br>A240a | 11 | 0.50 ± 0.01 | 0.20 ± 0.02 | 2500 ± 300 | 44 |
| **KE703b <sup>e</sup>** | S20a Y48F W72V G101M<br>S138Q H166N V168A<br>A204V A240a | 9 | N.D. | N.D. | 5500 ± 100 | 96 |

<sup>a</sup> Mutations are relative to the starting template (HG3 or KE70).
<sup>b</sup> Data represent the average of six to fifteen individual replicate measurements from two to five independent protein batches. N.D. indicates that individual parameters $K_M$ and $k_{cat}$ could not be determined accurately because saturation was not possible at the maximum substrate concentration tested (2 mM, Supplementary Figure 4 and Supplementary Figure 6), which is the substrate's solubility limit. In these cases, catalytic efficiencies ( $k_{cat}/K_M$ ) were calculated from the slope of the linear portion ( $[S] \ll K_M$ ) of the Michaelis-Menten model ( $v_0 = (k_{cat}/K_M)[E_0][S]$ ). Errors of regression fitting, which represent the absolute measure of the typical distance that each data point falls from the regression line, are provided.
<sup>c</sup> Values in parentheses are from Broom *et al.* <sup>14</sup> (HG series), or Röthlisberger *et al.* <sup>3</sup> and Khersonsky *et al.* <sup>10</sup> (KE70 series).
<sup>d</sup> HG649 is one of the 55 designs that passed all filtering criteria prior to the final molecular dynamics (MD) step.
<sup>e</sup> These enzymes contain two insertions, S20a and A240a/G240a.
<sup>f</sup> The KE702 design was inactive.

### Kinetic characterization of designs

All nine designs catalyzed the Kemp elimination (Supplementary Figure 4). The most active of these variants, HG630, had a catalytic efficiency of 15,000 ± 500 M^−1^ s^−1^, a value that is 120-fold higher than that of the starting template HG3 (Table 1). Furthermore, the three most active designs (HG198, HG259 and HG630) were more catalytically efficient than the HG3.3b variant obtained after three rounds of directed evolution and screening of >3000 mutants^8^. These designs all contain the M84C mutation that was previously shown to increase catalytic efficiency by one order of magnitude when combined with K50Q to yield the HG3-K50Q/M84C double mutant^15^ (Table 1). Analysis of mutant sequences suggested that introduction of the Q90F mutation into HG630 would further enhance its catalytic efficiency, as it doubles *k*_cat_/*K*_M_ when introduced into HG57 to yield HG44. Introduction of this mutation into HG630 resulted in a hexamutant (HG649) that is 256-fold more catalytically efficient than HG3 (*k*_cat_/*K*_M_ = 32,000 ± 3000 M^−1^ s^−1^) and more active than HG3.7, a variant found after seven rounds of directed evolution involving the screening of >7000 mutants^8^ (Table 1, Figure 2a). HG649 shares a majority of its mutation sites with HG3.7 but the combination of specific amino-acid substitutions is distinct, with only two mutations in common (Figure 2b). These results demonstrate how ensemble-based enzyme design produces similar activity increases as directed evolution but at a fraction of the experimental screening cost. Interestingly, HG649 was one of the 55 designs that passed all filtering criteria except for the final molecular dynamics analysis, which led to its elimination due to its low predicted catalytic competency (Supplementary Figure 3). This false negative, along with the two false positives (Table 1, HG378 and HG57), indicate that improvements are required to make the molecular dynamics filtering more robust and suggests that other eliminated sequences may also display high catalytic efficiency.

**Figure 2.**
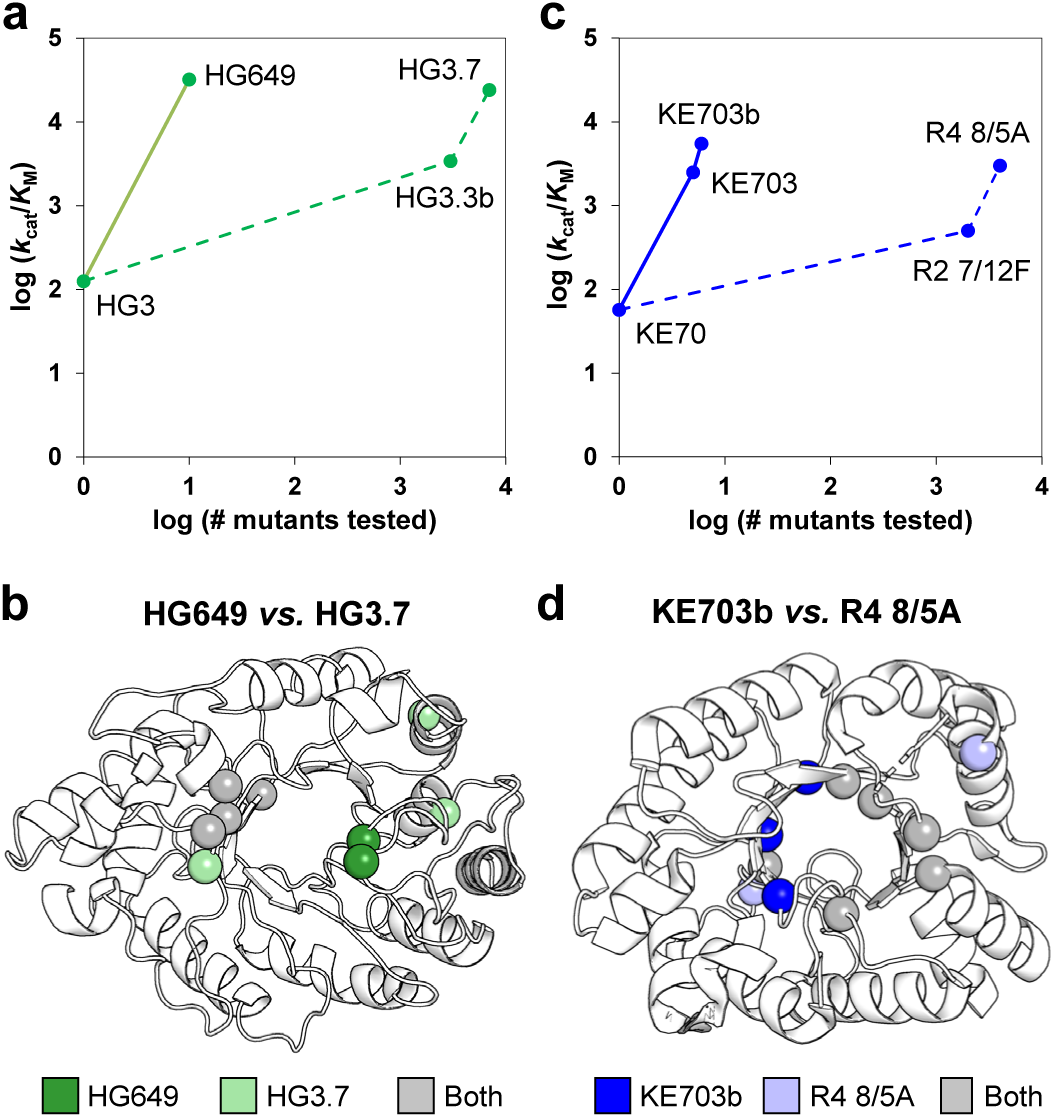
Ensemble-based design vs. directed evolution. (a) Ensemble-based design yields catalytic efficiency improvements comparable to those achieved by directed evolution with a fraction of the experimental screening effort. Full and dashed lines show ensemble-based design and directed evolution trajectories, respectively. (b) Ensemble-based design identifies a majority of the mutation sites (spheres) found by directed evolution (4/6) but mutation combinations of designed (HG649) and evolved (HG3.7) enzymes are distinct, as these share only two mutations (K50Q and M84C). (c) Full and dashed lines show ensemble-based design and directed evolution trajectories, respectively. KE703b differs from KE703 by reversion of its D16/N45 residues to the H16/D45 catalytic dyad found in KE70 and its evolved variants. (d) Designed variant KE703b shares 6 of the 9 mutation sites (spheres) found by directed evolution but has only three mutations in common with evolved variant R4 8/5A (S20a, Y48F, A204V).

### Crystal structures

To evaluate whether the catalytic efficiency of our designs was increased via the predicted molecular mechanisms (Figure 1b), we solved room-temperature (280 K) X-ray crystal structures of four of the most active designs (HG185, HG198, HG630 and HG649) in the presence and absence of the 6NT transition-state analogue. We used room-temperature X-ray crystallography for direct comparison with the HG3 non-cryogenic structure used to generate the ensemble and because this method can reveal conformational heterogeneity in protein structures that would not be visible at cryogenic temperatures, providing insights into the conformational ensemble that is sampled by a protein^21,24^. Crystals of HG185, HG630, and HG649 were obtained in the absence of 6NT, and soaked into solutions containing this molecule prior to X-ray data collection to obtain the analogue-bound structures (Supplementary Table 4). These three enzyme variants all crystallize in space group P 2_1_ with two molecules in the asymmetric unit, roughly matching the crystal form reported for HG3^14^. On the other hand, HG198 could only be crystallized in the presence of 6NT, and its unbound structure was derived from serially soaking these crystals in solutions lacking the transition-state analogue prior to data collection. In the presence of 6NT, HG198 crystallizes in the low symmetry space group P 1 with three molecules in the asymmetric unit, but removing the transition-state analogue via serial soaking transforms the crystals to space group P 2_1_ with a single molecule in the asymmetric unit. This unusual transformation of the crystal system was consistent over multiple crystal specimens. All crystals diffracted at resolutions of 1.32–1.90 Å, which enabled us to create high-quality structural models (Supplementary Table 5). All designs bound 6NT in the same catalytically productive pose (Figure 3a) as that observed in HG3, in excellent agreement with the computational models generated by ensemble-based design (Figure 4) but in stark contrast with those generated using the average crystal structure (Supplementary Figure 5). In this pose, the acidic N–H bond of 6NT that mimics the cleavable C–H bond of the substrate is located within hydrogen-bonding distance to the carboxylate oxygen of D127, while the basic nitrogen atom corresponding to the phenolic oxygen of the transition state forms a hydrogen bond with the N_ε_ atom of Q50, similar to what was observed for evolved variant HG3.7^14^. By contrast, HG3 does not form this second hydrogen bond because its K50 residue does not adopt a suitable side-chain conformation and instead it is a water molecule that hydrogen-bonds to the basic nitrogen atom of 6NT. In all cases, angles defining the hydrogen bond between D127 and the transition-state analogue were improved or within error compared to HG3 (Supplementary Table 3). In contrast to all other enzymes, the 6NT transition-state analogue of HG185 refined to an occupancy of approximately 60%, indicating that its crystal comprises a mixture of enzyme molecules in their bound and unbound forms. Given that the soaking procedure we used to generate crystals of HG185 bound to 6NT was identical to the one used to soak isomorphous crystals of HG630 and HG649 (Supplementary Table 4), which yielded 6NT bound at 100% occupancy, it is likely that the lower occupancy of this ligand in HG185 reflects its weaker binding in the active site, in agreement with the enzyme’s 6.5–14-fold lower catalytic efficiency compared to HG630 and HG649 (Table 1). The lower occupancy of 6NT in the HG185 structure is reflected by weaker electron density in the active site and the presence of an alternate D127 rotamer that does not hydrogen-bond with the transition-state analogue (Figure 3a). Regardless, the active site of all designed variants is more complementary to the transition state than that of HG3, as indicated by the lower solvent-accessible surface area for 6NT when bound in the active site of these variants (Figure 3b).

**Figure 3.**
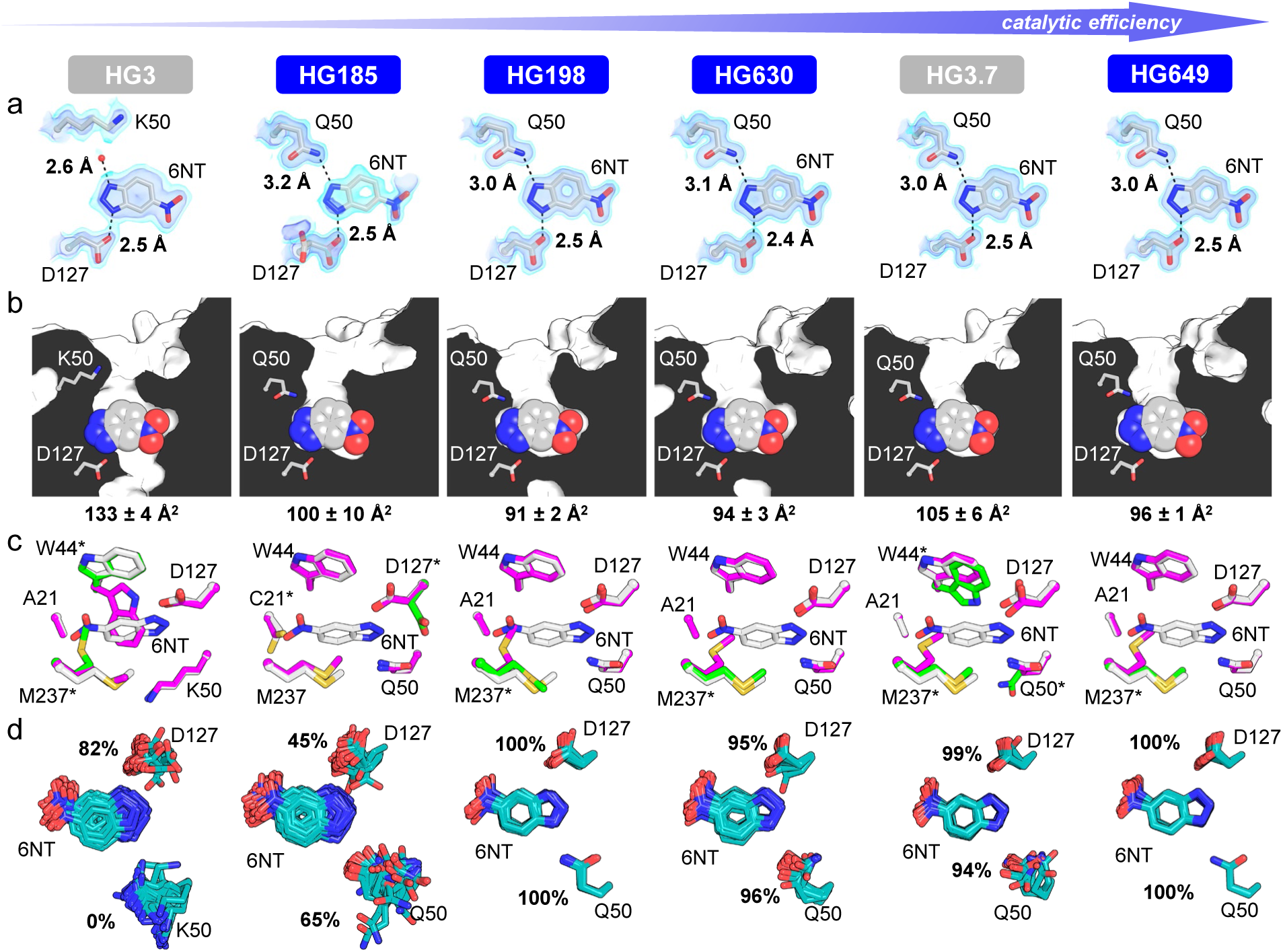
Crystal structures of HG-series Kemp eliminases. In all cases, only atoms from chain A are shown. Structures of HG3 (PDB ID: 5RG4 and 5RGA) and its evolved variant HG3.7 (PDB ID: 5RG6 and 5RGC) were previously published^14^. (a) Hydrogen bonds to the 6-benzotriazole (6NT) transition-state analogue are shown as dashed lines and distances are indicated. The red sphere represents a water molecule. The 2Fo-Fc map is shown in volume representation at two contour levels: 0.5 eÅ^−3^ and 1.5 eÅ^−3^ in light and dark blue, respectively. In the case of HG185, occupancy of 6NT in the crystal is approximately 60%, which is reflected by its lower electron density. (b) Cut-away view shows that active sites of designs are more complementary to the transition-state analogue than that of the HG3 template, as indicated by lower solvent-accessible surface areas for 6NT (mean ± s.d. across all chains). 6NT is shown as spheres. (c) Superposition of the 6NT-bound structure (white) with the highest (magenta) and lowest (green) occupancy conformers of the unbound structure for each Kemp eliminase. Active-site residues that are not preorganized are indicated by an asterisk. In the starting template HG3 and evolved variant HG3.7, the unbound state is never preorganized for catalysis as both W44 and M237 in the major and minor conformer, respectively, adopt side-chain rotamers that would prevent productive binding of the transition state. In the HG185 design, C21 is never preorganized in the unbound state, and the catalytic base D127 is only preorganized in the major conformer (80% occupancy). In the HG198, HG630 and HG649 designs, only M237 adopts a non-productive conformation in the unbound state, with an occupancy ranging from 47 to 64%. (d) Ensemble refinement of 6NT-bound structures show that designed variants HG198, HG630 and HG649, as well as evolved variant HG3.7, display lower structural heterogeneity than HG3 and a higher percentage of ensemble members whose catalytic side chains (D127 and Q50) make hydrogen bonds with 6NT. In the case of HG185, the lower occupancy of 6NT in the crystal (approximately 60%) likely contributes to its comparable heterogeneity to HG3 and lower percentage of ensemble members making catalytic contacts with 6NT.

**Figure 4.**
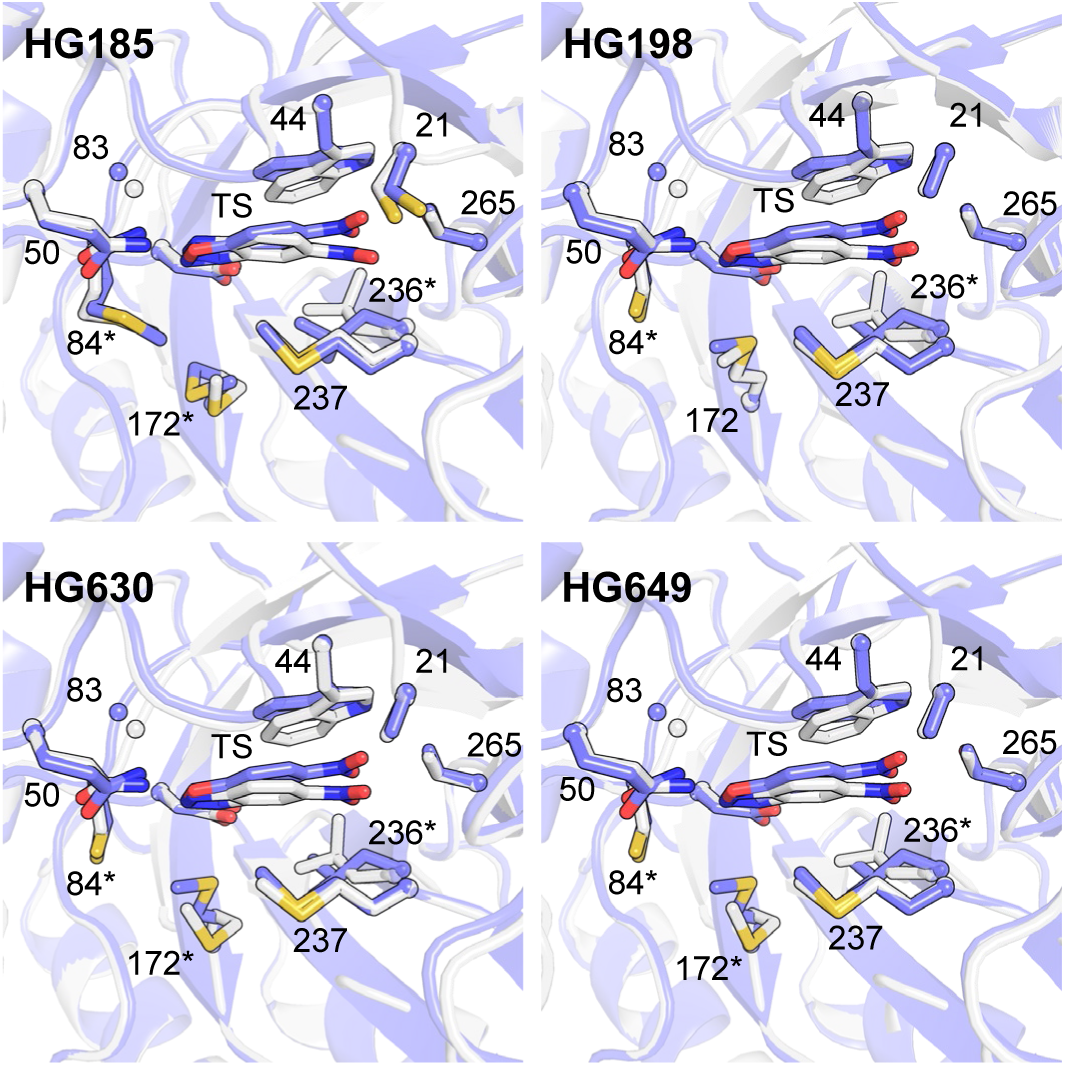
Ensemble-based design accurately predicts crystal structures of designed enzymes. Crystal structures of HG-series Kemp eliminases with bound 6NT (white) are overlaid on the corresponding design models obtained by ensemble-based design (blue). For clarity, only the major conformer of Chain A is shown. The transition state (TS) and transition-state analogue are shown at the center of each image. Side chains of all residues forming the binding pocket are shown with the exception of P45, which was omitted for clarity. The sphere shows the alpha carbon of G83. Asterisks indicate residues that adopt side-chain rotamers varying by >20 degrees around one or more side-chain dihedrals between the design model and crystal structure.

Next, we evaluated active-site preorganization by comparing structures in the presence and absence of bound 6NT. In HG3, the unbound form is never preorganized for catalysis as both W44 and M237 adopt conformations that would prevent productive binding of 6NT in the major or minor conformations, respectively (Figure 3c). This is also the case for evolved variant HG3.7^14^. HG185 is also never preorganized in the unbound form because C21 exclusively adopts a rotamer that would clash with 6NT. Furthermore, the D127 catalytic residue of HG185 adopts a low-occupancy (7% and 14% for chains A and B, respectively), catalytically non-productive conformation in the unbound form that cannot interact favorably with 6NT, likely contributing to its inferior activity compared with the other variants. In contrast, active sites of higher activity variants HG198, HG630 and HG649 are correctly preorganized for catalysis in a substantial portion of the molecules in the crystal, with only M237 adopting a non-productive conformation at 47–64% occupancy (Figure 3c). Overall, crystal structures confirmed that activity of designed HG variants was increased by several of the mechanisms known to enhance enzyme catalysis and that we aimed to reproduce using our filtering criteria.

### Enrichment of catalytically competent substates

The ability of an enzyme to frequently sample productive substates in its conformational ensemble is another important feature of efficient catalysis^15,17,25,26^. To evaluate this property for our designs, we performed ensemble refinement of their 6NT-bound structures and evaluated the proportion of ensemble members that make the designed catalytic contacts with the transition-state analogue. We observed that in HG3, the hydrogen bond between the catalytic base and 6NT is formed in 82% of ensemble members, and no interaction is observed between the ligand and K50 (Figure 3d). In HG185, interactions between 6NT and catalytic residues D127 and Q50 are present in 45% and 65% of ensemble members, respectively, likely due to the lower occupancy of 6NT. In the case of the most active variants HG198, HG630 and HG649, catalytic contacts to both D127 and Q50 are present in ≥95% of ensemble members, and their ensembles are less structurally diverse than those of HG3 or HG185 (Figure 3d, Figure 5a, Supplementary Table 6). The higher percentage of hydrogen bonds between 6NT and catalytic residues and narrower conformational ensembles of high activity HG variants whose structures contain transition-state analogues at 100% occupancy confirm a higher population of catalytically-competent substates than in the ensemble of HG3.

**Figure 5.**
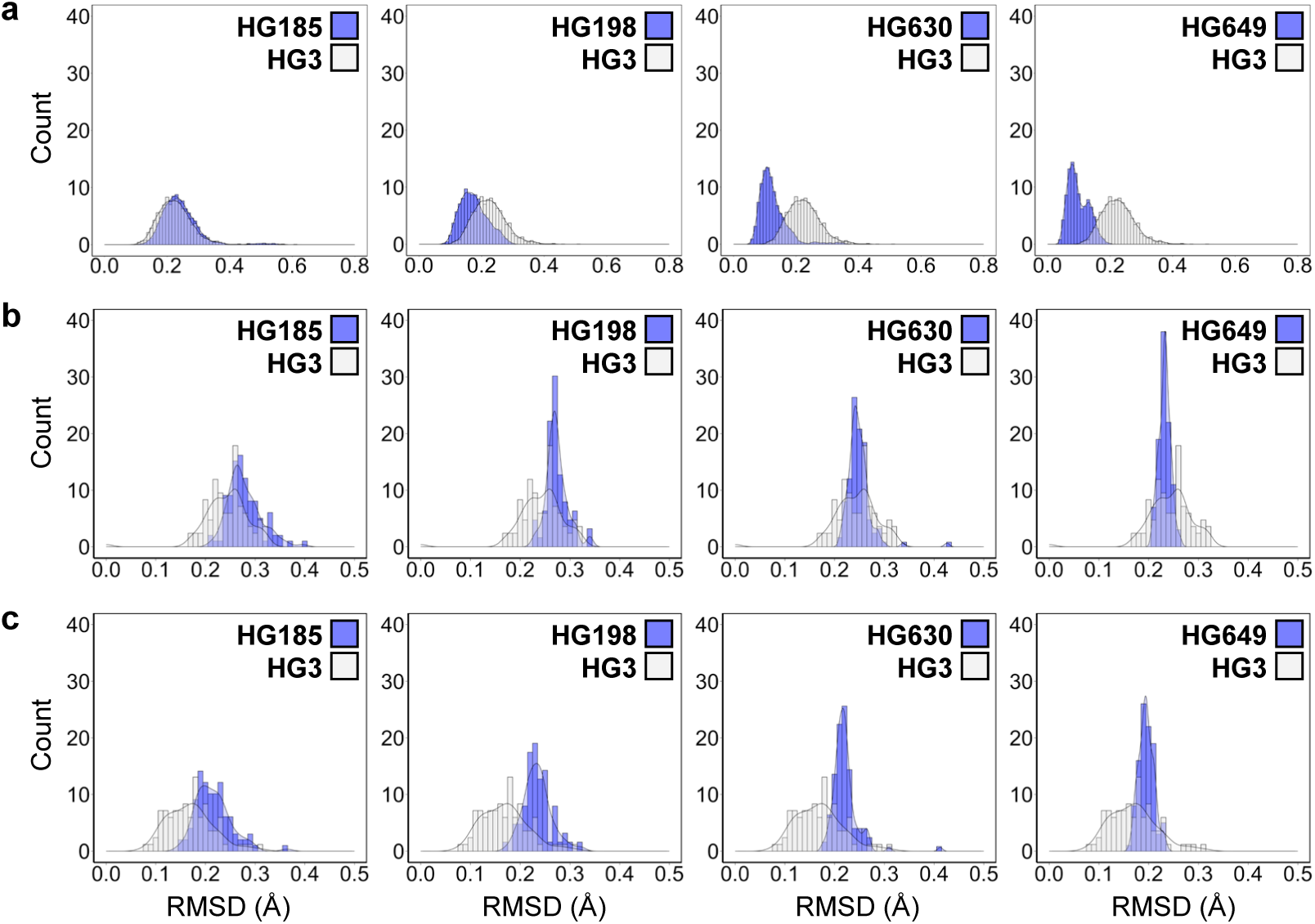
Enrichment of catalytically-competent substates in the conformational ensemble. Distributions of pairwise backbone root-mean-square deviations (RMSD) for designed active-site residues (listed in Supplementary Table 7) calculated between each backbone template obtained by ensemble refinement of various HG-series Kemp eliminases with bound 6NT and (a) each template within the same ensemble, (b) the bb05 template used for computational design, and (c) the HG3 crystal structure with bound 6NT (PDB ID: 5RGA^14^). Distributions show that (a) with the exception of HG185, ensemble members are more similar to each other in the ensembles of higher activity variants than in the ensemble of HG3, (b) the bb05 template is equally or more similar to the ensembles of the highest activity variants HG630 and HG649 (blue) than to its own ensemble (white), whereas (c) the HG3 crystal structure is more similar to its own ensemble (white) than to those from higher activity HG variants (blue). Median RMSD values and Z-scores for comparison of distributions are reported on Supplementary Table 6.

Having established that the conformational ensembles of efficient HG variants were enriched in catalytically competent substates, we investigated whether the bb05 template that we extracted from the HG3 ensemble and used as design template is a better approximation of the transition-state ensemble than the HG3 crystal structure obtained by traditional refinement of a single, major conformation. To do so, we calculated the pairwise backbone root-mean-square deviations of designed active-site residues (Supplementary Table 7) between every member of the HG variants’ ensembles and bb05. Distributions of root-mean-square deviations show that the bb05 active-site backbone is as or more structurally similar to the ensembles of the two most active variants (HG630 and HG649) than to its own ensemble (Figure 5b, Supplementary Table 6). By contrast, the HG3 crystal structure with bound 6NT is more similar to its own ensemble than to those of all higher activity HG variants (Figure 5c, Supplementary Table 6). These results strongly support our hypothesis that ensemble refinement performed on the crystal structure of a low activity *de novo* enzyme allows sampling of catalytically-productive conformational substates for more accurate accommodation of the transition state than the single crystal structure.

### KE70 designs

To test the general applicability of our ensemble-based enzyme design method, we applied it to another Kemp eliminase that was designed on a distinct protein scaffold. This enzyme, KE70, contains a His/Asp dyad that acts as catalytic base and a Ser hydrogen-bond donor to stabilize the phenolic oxygen of the transition state^3^. To allow direct comparison to our HG variants, we designed variants using a theozyme comprising an Asp residue as catalytic base and a Gln as hydrogen-bond donor. Starting from the available unbound cryogenic crystal structure of KE70 (PDB ID: 3NPV^10^), we generated an 84-member ensemble (Supplementary Table 1) and used it to design variants predicted to efficiently catalyze the Kemp elimination using a similar protocol as that used for HG variants but without the final filtering step involving molecular dynamics (Methods). Three variants were selected for kinetic analysis, and these contain between 11 and 13 mutations (Supplementary Table 8), including the D16 and Q138 catalytic motif as well as two insertions in active site loops (S20a and A240a) that we introduced after design to allow for comparison with evolved KE70 variants that contain these insertions^10^. Two of the three tested designs were active (Table 1, Supplementary Figure 6). The most active variant, KE703, displays a *k*_cat_/*K*_M_ of 2500 ± 300 M^−1^ s^−1^, which is 44-fold higher than KE70 and approximately 17-fold higher than that of a KE70 variant containing the two loop insertions (Table 1). To compare with evolved KE70 variants that contain the His/Asp dyad as catalytic base, we introduced the D16H and N45D mutations into KE703, generating the KE703b variant. This variant has a *k*_cat_/*K*_M_ of 5500 ± 100 M^−1^ s^−1^, which is approximately 100-fold higher than KE70 and two-fold higher than that of the evolved variant R4 8/5A that was isolated after 4 rounds of directed evolution and screening of approximately 4000 clones^10^ (Figure 2c). KE703b shares a majority of its mutation sites with R4 8/5A but its specific amino-acid substitutions are distinct, with only three mutations in common (Figure 2d). These results confirm the general applicability of ensemble-based design and its ability to generate artificial enzymes that are orders of magnitude more active than the original *de novo* enzymes at a fraction of the experimental effort required by directed evolution. Furthermore, our results show that ensemble-based design does not require an ensemble generated from a crystal structure with a bound transition-state analogue to be successful, which is advantageous given that such structures are less common than unbound ones.

## Discussion

In this work, we used ensemble-based design to increase the catalytic efficiency of the HG3 and KE70 *de novo* Kemp eliminases by two orders of magnitude, yielding artificial enzymes with *k*_cat_/*K*_M_ values that are within the 10^3^–10^6^ M^−1^ s^−1^ range encompassing a majority of natural enzymes^27^, and only 4- or 10-fold lower than those of the most active variants evolved from these enzyme prototypes (e.g., *k*_cat_/*K*_M_ of 126,000 M^−1^ s^−1^ and 53,100 M^−1^ s^−1^ for HG3.17^14^ and KE70 R8 12/12B^10^, respectively). To achieve these results, we tested a number of variants that is several orders of magnitude lower than those previously used to achieve comparable improvements by directed evolution^8,10^. Further increases to catalytic efficiency could be achieved by incorporating beneficial mutations outside the active site, which have been shown to augment catalysis via alterations to allosteric networks^28^. Although more challenging to predict a priori, mutational hotspots distal from the active site can be identified using nuclear magnetic resonance spectroscopy^29^ or by applying correlation-based tools to molecular dynamics trajectories^30^. In theory, it should be possible to create artificial enzymes for any desired reaction with catalytic efficiencies comparable to that of the average natural enzyme^27^ using purely rational approaches, and the ensemble-based design procedure described here can expedite this process.

The success of our ensemble-based design relied on exploiting the spatiotemporally-averaged information contained in X-ray crystallographic data to overcome the backbone sampling challenge that is responsible for much inaccuracy in enzyme design^8^. While others have previously attempted to address the backbone sampling issue by using ensembles generated via Monte-Carlo-based backrub sampling and incorporating loop remodelling during the enzyme design process, activity improvements were modest (<5-fold)^10^. These results, in combination with our previous observation that use of an ensemble generated by unconstrained molecular dynamics does not allow accurate recapitulation of catalytic contact and active-site configurations for an efficient artificial enzyme^14^, suggest that to obtain large activity increases, it is not sufficient to introduce backbone flexibility in the design process. Instead, the correct backbone conformation must be sampled. Crystallographic ensemble refinement of the initial *de novo* enzyme satisfies this stringent requirement in that the resulting model represents the true conformational ensemble sampled by the enzyme, which should include productive substates that are better suited for stabilization of the transition state than the average crystal structure. Generating the ensemble from a structure with a bound transition-state analogue likely helps to enrich such productive substates, and could partially explain why the KE70 ensemble, which was derived from an unbound structure, led to a lower overall activity improvement. Although the use of an ensemble derived from enzymes in their bound and unbound forms led to activity improvements, one important disadvantage of our approach is that it requires a crystal structure of the original *de novo* enzyme to generate the ensemble. Since *de novo* enzymes are typically generated from small, monomeric protein scaffolds whose crystal structures have been determined to high resolution, we do not anticipate that structure determination will create a substantial bottleneck in most cases.

Our results support the idea that to design efficient artificial enzymes, it is essential to consider residual conformational heterogeneity present in the designs, and whether it leads to the population of catalytically unproductive conformations. In other words, dynamic aspects of enzyme structure, such as active-site preorganization and enrichment of catalytically-competent substates, must also be considered. This is supported by the fact that the most active designs described here, HG630 and HG649, were ranked lower than less active ones such as HG44 and HG57 based on their predicted ability to stabilize the transition state (as reflected by computed energies, Supplementary Table 2), which has hitherto been the most important metric used to pick designs for experimental testing. Recently, researchers have proposed that enzyme design should be viewed as a population shift problem, where relative stabilities of conformational substates are redistributed by the introduction of mutations to stabilize productive substates^15,28^. Previous work on the HG series of Kemp eliminases by our team^14^ and others^15^ demonstrated that directed evolution improves activity through essentially the same mechanism, with higher activity variants having reduced flexibility and greater active-site preorganization, both of which favor the population of catalytically active substates. The similarity is further highlighted by the fact that our ensemble-based design approach and directed evolution identify activity enhancing mutations at many of the same positions.

Moving forward, it should be possible to further enhance the population of catalytically-competent conformations for a designed enzyme ensemble using the information obtained from ensemble refinement of X-ray diffraction data. Specifically, we anticipate using these ensembles to identify sequences that can stabilize productive substates while simultaneously destabilizing non-productive ones in a multistate design approach^31^. The ability to remodel an enzyme conformational landscape^32^ in this way, using negative design strategies to widen the energy gap between productive and non-productive conformations, may open the door to the design of more efficient artificial enzymes than previously possible. Beyond the scope of enzyme design, we expect that a similar strategy, which takes advantage of crystallographically-derived ensemble models, could find utility in a broad range of protein design applications. For example, directed evolution following computational design of protein-protein interactions has been shown to enhance affinity by eliminating conformations from the ensemble that are non-productive for binding^24^. Additionally, conformational heterogeneity has been shown to be detrimental to the functions of computationally designed fluorescent proteins^33^ and ligand-binding proteins^34^. Each of these observations suggests a potential application of our proposed approach.

## Methods

### Ensemble generation

Ensembles of backbone templates were generated using the ensemble refinement protocol^18^ implemented in the PHENIX macromolecular structure determination software (v1.18.2-3874)^35^. In ensemble refinement, local atomic fluctuations are sampled using molecular dynamics simulations accelerated and restrained by electron density to produce ensemble models fitted to diffraction data. Briefly, input crystal structures were edited to remove low-occupancy conformers and assign an occupancy of 1.0 to the remaining conformer. Following addition of riding hydrogens, parallel ensemble refinement simulations were performed using various combinations of the parameters p_TLS_ (0.6, 0.8, 0.9, 1.0), τ_x_ (0.5, 1.5, 2.0) and w_x-ray_ (2.5, 5.0, 10.0), where p_TLS_ describes the percentage of atoms included in a translation-libration-screw (TLS) model use to remove the effects of global disorder, τ_x_ is the simulation time-step and w_x-ray_ is the coupled tbath offset, which controls the extent to which the electron density contributes to the simulation force field such that the simulation runs at a target temperature of 300 K. The ensemble generated from each crystal structure that displayed the lowest R_free_ value was selected for design and analysis (Supplementary Table 1).

### Computational enzyme design

All calculations were performed with the Triad protein design software^36^ (Protabit, Pasadena, CA, USA) using a Monte Carlo with simulated annealing search algorithm for rotamer optimization. Individual backbone templates from each structural ensemble were prepared for computational design using the *addH.py* application within Triad, which optimized and renamed hydrogen atoms, assigned protonation state to His residues, flipped His/Asn/Gln sidechains if necessary, and rotated hydroxyl, sulfhydryl, methyl, and amino groups in Ser/Thr, Cys, Met, and Lys, respectively.

The Kemp elimination transition-state (TS) was built using parameters described by Privett and coworkers^7^. For HG3 designs, D127 was used as the catalytic base, and Gln/Asn/Thr/ Ser were tested as hydrogen-bond donors at various positions (Supplementary Table 7). For KE70 designs, D16 served as the catalytic base (corresponding to position 17 in the KE70 crystal structure, PDB ID: 3NPV), Q138 as hydrogen-bond donor, and F48 was utilized for introduction of a pi-stacking interaction with the TS (Supplementary Table 9). For both HG3 and KE70 designs, all other positions in the active site were mutated to Gly during theozyme placement to avoid steric clashes with designed catalytic residues. The 2002 Dunbrack backbone-independent rotamer library^37^ with expansions of ± 1 standard deviation around χ_1_ and χ_2_ was used to provide side-chain conformations. A library of TS poses was generated in the active site by targeted ligand placement^1^ using the contact geometries listed in Supplementary Table 10. TS pose energies were calculated using a modified version of the Phoenix protein design energy function^36^ consisting of a Lennard-Jones 12–6 van der Waals term from the Dreiding II force field^38^ with atomic radii scaled by 0.9, a direction-dependent hydrogen bond term with a well depth of 8.0 kcal mol^−1^ and an equilibrium donor-acceptor distance of 2.8 Å^39^, and an electrostatic energy term modeled using Coulomb’s law with a distance-dependent dielectric of 10. During the energy calculation step, TS–side-chain interaction energies were biased to favor interactions that satisfy contact geometries (Supplementary Table 11) by applying an energy benefit of –100 kcal mol^−1^ as described by Lassila *et al.*^1^. A Triad patch file for theozyme placement is provided as a Supplementary File.

Next, active-site repacking calculations were performed on the theozyme-containing templates. In these calculations, the TS was translated ± 0.4 Å in each Cartesian coordinate in 0.2-Å increments and rotated 10° about all three axes (origin at TS geometric center) in 5° increments for a total combinatorial rotation/translation search space of 15,625 possible poses. The identities of catalytic residues were fixed and allowed to sample all rotamers of that amino-acid type. Residues that were converted to Gly in the theozyme placement step were allowed to sample rotamers of selected amino acids (Supplementary Tables 7 and 9) to form a mostly hydrophobic active site. Side-chain–TS interaction energies were biased to favor contacts satisfying geometries as done during the theozyme placement step (Supplementary Table 11). Rotamer optimization was carried out using the search algorithm, rotamer library, and energy function described above. A Triad patch file for active-site repacking is provided as a Supplementary File.

After active-site repacking, we calculated energy differences between each repacked structure and the corresponding structure in the absence of the TS where all designed residues (including catalytic residues) were converted to Gly. This energy subtraction was done to allow comparison of energies for sequences across multiple backbone templates. We also measured angles defining the hydrogen-bond contact between the catalytic base and the TS of the top-scoring sequence obtained from each backbone template. Sequences from the HG3 and KE70 backbone template that gave the lowest energy top-ranked sequence with angles between its catalytic base and the transition state within 3% or 3.5% error of the optimal geometries calculated for acetamide dimers^23^ (Supplementary Figure 2, Supplementary Table 3), respectively, were selected for library design and cleaning, as described below.

### Computational library design

Library design was performed with the CLEARSS algorithm^40^ using the energies obtained after active-site repacking on the single template selected as described above. For a specific library size configuration, which is the specific number of amino acids at each position in the protein, the highest probability set of amino acids at each position (based on the sum of their Boltzmann weights) were included in the library. A combinatorial library of approximately 1000 sequences derived from the selected HG3 or KE70 ensemble member was generated. To ensure that the lowest energy rotameric configuration was obtained for each library sequence, we used the *cleanSequences.py* app in Triad to “clean” their structures by optimizing rotamers on the input backbone template using the active-site repacking protocol described above but without modification of catalytic residue rotameric configuration. To compare energies of “cleaned” sequences, we calculated the energy difference between each optimized structure and the corresponding structure in the absence of the TS where all designed residues (including catalytic residues) were converted to Gly. These energies are reported throughout the figures and text.

### Design filtering

To decrease the number of sequences to test experimentally, “cleaned” structures were filtered using geometrical criteria. We kept sequences with (i) a TS solvent-accessible surface area lower than 100 Å^2^ or 102 Å^2^ for HG3 and KE70 designs, respectively, and (ii) ≥70 % of all designed active-site residues predicted to be preorganized, which was determined as those that adopted the same rotamer in the absence and presence of TS. Furthermore, sequences that contained ≥5 Met residues or ≥2 Cys residues were discarded to reduce conformational heterogeneity of the active site and prevent formation of disulfide bonds, respectively. A final filtering step consisting of short molecular dynamics simulations was performed for HG3 designs, as described below.

### Unconstrained molecular dynamics simulation for filtering of HG3 designs

All simulations were performed using the Amber 2020 software (http://ambermd.org/) with the AMBER19SB forcefield^41^. Long-range electrostatics (>10 Å) were modeled using the particle mesh Ewald method^42^, and a time step of 2 fs was used for the production phase. Parameters for the 5-nitrobenzisoxazole substrate were generated using the Antechamber^43^ package. “Cleaned” computational models generated by Triad for the 55 sequences that passed the filtering procedure described above were prepared for molecular dynamics. The TS structure was converted to that of the substrate, hydrogen atoms were added using Reduce^44^, and each substrate-bound protein molecule was placed in a dodecahedral box with periodic boundary conditions where the distance between the protein surface and the box edges were set to 10 Å. After explicit TIP3P water molecules^45^ were added, charges on protein atoms were neutralized with Na^+^ and Cl^−^ counter-ions at a concentration of 0.15 M. The structures were then energy minimized with the steepest descent method to a target maximum force of 1000 kJ mol^−1^ nm^−1^. Before equilibration, the system was heated to a target temperature of 300 K for 100 ps. The system was then equilibrated under an NVT ensemble for 100 ps at a temperature of 300 K using a Nose-Hoover thermostat^46^, while applying position restraints for heavy protein atoms. A second equilibration step under an NPT ensemble was performed for 100 ps with constant pressure and temperature of 1 bar and 300 K, respectively, using the Berendsen barostat^47^. Following removal of position restraints, 20-ns production runs under Parrinello-Rahman pressure coupling^48^ were initiated from the final snapshot of the NPT equilibration. After molecular dynamics, designs were evaluated and ranked according to their catalytic competency, which is the percentage of snapshots in which catalytic contacts between substrate and catalytic side chains were maintained, as calculated using distance and angle metrics listed on Supplementary Figure 3.

### Protein expression and purification

Codon-optimized (*E. coli*) and his-tagged (C-terminus) genes for Kemp eliminases (Supplementary Tables 12–13, Supplementary Figures 7–8) cloned into the pET-29b(+) vector via *Nde*I and *Xho*I were obtained from Twist Bioscience. Enzymes were expressed in *E. coli* BL21-Gold (DE3) cells (Agilent) using lysogeny broth (LB) supplemented with 100 μg mL^−1^ kanamycin. Cultures were grown at 37 °C with shaking to an optical density at 600 nm of 0.3–0.7, and protein expression was initiated with 1 mM isopropyl-β-D-1-thiogalactopyranoside. Following incubation at 16 °C for 16 h with shaking (250 rpm), cells were harvested by centrifugation, resuspended in 8 mL lysis buffer (5 mM imidazole in 100 mM potassium phosphate buffer pH 8.0) supplemented with 1.25 g mL^−1^ of lyophilized lysozyme and 1 U mL^−1^ of benzonase nuclease (Merck Millipore), and lysed with an EmulsiFlex-B15 cell disruptor (Avestin). Proteins were purified by immobilized metal affinity chromatography using Ni–NTA agarose (Qiagen) pre-equilibrated with lysis buffer in individual Econo-Pac gravity-flow columns (Bio-Rad). Columns were washed twice, first with 10 mM imidazole in 100 mM potassium phosphate buffer (pH 8.0), and then with the same buffer containing 20 mM imidazole. Bound proteins were eluted with 250 mM imidazole in 100 mM potassium phosphate buffer (pH 8.0) and exchanged into 100 mM sodium phosphate buffer (pH 7.0) supplemented with 100 mM sodium chloride using Econo-Pac 10DG desalting pre-packed gravity-flow columns (Bio-Rad). Proteins used for crystallization were further purified by gel filtration in 50 mM sodium citrate buffer (pH 5.5) and 150 mM sodium chloride using an ENrich SEC 650 size-exclusion chromatography column (Bio-Rad). Purified samples were concentrated using Amicon Ultracel-3K centrifugal filter units (EMD Millipore) and quantified by measuring the absorbance of the denatured protein in 6 M guanidinium chloride at 280 nm and applying Beer-Lambert’s law using calculated extinction coefficients obtained from the ExPAsy ProtParam tool (https://web.expasy.org/protparam).

### Steady-state kinetics

All assays were carried out at 27 °C in 100 mM sodium phosphate buffer (pH 7.0) supplemented with 100 mM sodium chloride. Triplicate 200-μL reactions with varying concentrations of freshly prepared 5-nitrobenzisoxazole (Aablocks) dissolved in methanol (10% final concentration, pH of reaction mixture adjusted to 7.0 after addition of methanol-solubilized substrate) were initiated by the addition of 0.02–5.0 µM or 0.03–16.0 µM for HG or KE70 variants, respectively. Product formation was monitored spectrophotometrically at 380 nm (ε = 15,800 M^−1^ cm^−1^) in individual 96-well plates (Greiner Bio-One) wells using a Biotek Synergy H1 multimode plate reader (Agilent). Path lengths for each well were calculated ratiometrically using the difference in absorbance of 100 mM sodium phosphate buffer (pH 7.0) supplemented with 100 mM sodium chloride and 10% methanol at 900 and 975 nm (27 °C). Linear phases of the kinetic traces were used to measure initial reaction rates. For cases where saturation was not possible at the maximum substrate concentration tested (2 mM), data were fitted to the linear portion of the Michaelis-Menten model (v_0_ = (*k*_cat_/*K*_M_)[E_0_][S]) and *k*_cat_/*K*_M_ was deduced from the slope. In all other cases, data were fitted to the Michaelis-Menten equation to calculate individual *k*_cat_ and *K*_M_ parameters.

### Crystallization

Enzyme variants were prepared in 50 mM sodium citrate buffer (pH 5.5) at the concentrations listed in Supplementary Table 4. For samples that were co-crystallized with the transition-state analogue, a 100 mM stock solution of 6-nitrobenzotriazole (6NT, AstaTech) was prepared in dimethyl sulfoxide (DMSO) and diluted 20-fold in the enzyme solutions for a final concentration of 5 mM (5% DMSO). For each enzyme variant, we carried out initial crystallization trials in 15-well hanging drop format using EasyXtal crystallization plates (Qiagen) and a crystallization screen that was designed to explore the chemical space around the crystallization conditions reported by Broom *et al*.^14^ Crystallization drops were prepared by mixing 1 μL of protein solution with 1 μL of the mother liquor and sealing the drop inside a reservoir containing an additional 500 μL of the mother liquor solution. This procedure was successful for HG185, HG198, and HG630; however, crystallization of HG649 required identification of a new crystallization condition using sparse matrix screening. We used commercial screening kits (NeXtal) to prepare crystallization drops in 96-well sitting drop plates by mixing 1 μL of protein solution with 1 μL of the mother liquor and sealing the drop inside a reservoir containing an additional 100 μL of the mother liquor solution. After identifying a suitable mother liquor solution from screening experiments, large crystals suitable for X-ray diffraction experiments were obtained in 15-well hanging-drop format as described for the other HG variants. The specific growth conditions that yielded the crystals of each HG variant used for X-ray data collection are provided in Supplementary Table 4. HG185, HG630, and HG649 crystallized in the absence of the 6NT transition-state analogue, and the resulting crystals were isomorphous with the HG3 crystal form reported in the PDB (PDB ID: 5RGA and 5RG4). For these variants, crystals with bound 6NT were obtained by soaking the unbound crystals in mother liquor containing 10 mM 6NT for at least 30 minutes prior to data collection. On the other hand, HG198 could not be crystallized without 6NT present, and the crystals that were obtained belonged to space group P1 with three polypeptides in the asymmetric unit. To prepare crystals of HG198 without 6NT bound, we serially soaked 6NT-bound crystals in three drops of crystallization mother liquor without 6NT. Each sequential soaking step was carried out for at least 30 minutes. After removal of 6NT by soaking, the P1 crystals undergo a transition to space group P2_1_, with a smaller unit cell and only one molecule per asymmetric unit. Although we could not determine the cause of this unusual change in crystal symmetry upon soaking, we observed that it occurred with high reproducibility. We measured 10 individual crystals of 6NT-bound HG198, which all belonged to space group P1 and were roughly isomorphous. Additionally, we measured four individual crystals of HG198 without 6NT, and found that all of them demonstrated the transition to monoclinic symmetry.

### X-ray data collection and processing

Prior to X-ray data collection, crystals were mounted in polyimide loops and sealed using a MicroRT tubing kit (MiTeGen). Single-crystal X-ray diffraction data were collected on beamline 8.3.1 at the Advanced Light Source. The beamline was equipped with a Pilatus3 S 6M detector, and was operated at a photon energy of 11111 eV. Crystals were maintained at 280 K throughout the course of data collection. Each data set was collected using a total X-ray dose of 50 kGy or less, and covered a 180° wedge of reciprocal space. Multiple data sets were collected for each enzyme variant either from different crystals, or if their size permitted, from non-overlapping volumes of larger crystals, in order to identify optimally diffracting samples. X-ray data were processed with the Xia2 0.5.492 program (https://doi.org/10.1107/S0021889809045701)^49^, which performed indexing, integration, and scaling with DIALS^50^, followed by merging with Pointless^51^. For each data set, a resolution cutoff was taken where the CC1/2 and <I/σI> values for the merged intensities fell to approximately 0.5 and 1.0, respectively. Information regarding data collection and processing is presented in Supplementary Table 5. The reduced diffraction data were analyzed with phenix.xtriage to check for crystal pathologies, and no complications were identified.

### Structure determination

We obtained initial phase information for calculation of electron density maps by molecular replacement using the program Phaser^52^, as implemented in v1.13.2998 of the PHENIX suite^53^. Several different models of HG variants, including crystal structures and design models, were used for molecular replacement. For all data sets except that of HG630 (+) 6NT, we performed iterative steps of manual model rebuilding followed by refinement of atomic positions, atomic displacement parameters, and occupancies using a translation-libration-screw (TLS) model, a riding hydrogen model, and automatic weight optimization. For HG630 (+) 6NT, refinement was done as described above but without usage of a translation-libration-screw (TLS) model. All model building was performed using Coot 0.8.9.236^54^ and refinement steps were performed with phenix.refine within the PHENIX suite (v1.13-2998)^53,55^. Restraints for 6NT were generated using phenix.elbow^56^, starting from coordinates available in the Protein Data Bank (PDB ligand ID: 6NT). Further information regarding model building and refinement, as well as PDB accession codes for the final models, are presented in Supplementary Table 5.

## Data availability

Structure coordinates for all HG-series Kemp eliminases have been deposited in the RCSB Protein Data Bank with the following accession codes: HG185 (8USK, 8USL), HG198 (8USI, 8USJ), HG630 (8USG, 8USH), HG649 (8USE, 8USF). Source data are provided with this paper. Other relevant data are available from the corresponding authors upon reasonable request.

## Code availability

Triad patch files are provided with this paper. Requests for the Triad protein design software should be addressed to Protabit (https://www.protabit.com/).

## Acknowledgements

R.A.C. acknowledges grants from the Natural Sciences and Engineering Research Council of Canada (RGPIN-2021-03484 and RGPAS-2021-00017) and the Canada Foundation for Innovation (26503). R.A.C. and M.C.T. acknowledge a joint grant from the Human Frontier Science Program (RGP0004/2022). This research was enabled in part by support provided by Compute Ontario (www.computeontario.ca) and the Digital Research Alliance of Canada (alliance can.ca). Beamline 8.3.1 at the Advanced Light Source is operated by the University of California San Francisco with generous support from the National Institutes of Health (R01 GM124149 for technology development and P30 GM124169 for user support), and the Integrated Diffraction Analysis Technologies program of the US Department of Energy Office of Biological and Environmental Research. The Advanced Light Source (Berkeley, CA) is a national user facility operated by Lawrence Berkeley National Laboratory on behalf of the US Department of Energy under contract number DE-AC02-05CH11231, Office of Basic Energy Sciences. The contents of this publication are solely the responsibility of the authors and do not necessarily represent the official views of NIGMS or NIH. The authors wish to thank Natalie K. Goto for critical reading of the manuscript.

## Author Contributions

R.V.R., M.C.T. and R.A.C. conceived the project. R.V.R. and N.Z. performed computational design experiments. R.V.R. and J.D.M.M. purified proteins. R.V.R. and J.D.M.M. performed enzyme kinetics experiments. B.S. crystallized proteins. J.M.R. and M.C.T. performed X-ray diffraction experiments. B.S., R.V.R., and R.A.C. performed refinements. M.C.T. designed X-ray crystallography experiments. R.A.C. and R.V.R. wrote the manuscript. N.Z., B.S., and M.C.T. edited the manuscript.

## Competing Interests

The authors declare no competing interests.

## Notes

### Competing Interest Statement

The authors have declared no competing interest.

