## Supplementary Information for "Design of efficient artificial enzymes using crystallographically-enhanced conformational sampling"

**This file includes:**

Supplementary Tables 1–13

Supplementary Figures 1–8

**Supplementary Table 1.** Ensemble refinement results

| Structure | PDB ID | Resolution (Å) | # chains | $R_{work}/R_{free}$ before ensemble refinement | $R_{work}/R_{free}$ after ensemble refinement | Ensemble size |
|---|---|---|---|---|---|---|
| **HG3 (+) 6NT** | 5RGA | 1.86 | 2 | 0.1482/0.1863 | 0.1332/0.1814 | 84 |
| **HG185 (+) 6NT** | 8USL | 1.55 | 2 | 0.1610/0.1831 | 0.1387/0.1740 | 100 |
| **HG198 (+) 6NT** | 8USJ | 1.60 | 3 | 0.1540/0.1900 | 0.1402/0.1890 | 50 |
| **HG630 (+) 6NT** | 8USH | 1.32 | 2 | 0.1190/0.1476 | 0.1316/0.1589 | 127 |
| **HG649 (+) 6NT** | 8USF | 1.45 | 2 | 0.1373/0.1567 | 0.1302/0.1584 | 102 |
| **KE70 (–) 6NT** | 3NPV | 1.48 | 2 | 0.1430/0.1700 | 0.1290/0.1549 | 84 |

**Supplementary Table 2.** Design filtering – HG series

| Design | Mutations from HG3 | Energy (kcal mol$^{-1}$) | SASA (Å$^2$) [a] | # Cys | # Met | # Preorganized residues (on 29) [b] | Residues not preorganized |
|---|---|---|---|---|---|---|---|
| **HG44** | A21C K50Q Q90F A125V | –206.8 | 86.2 | 1 | 4 | 23 | Cys21, Trp44, Gln50, Val125, Leu236, Asn239 |
| **HG57** | A21C K50Q Q90L A125V | –205.9 | 86.2 | 1 | 4 | 21 | Cys21, Trp44, Gln50, Val125, Leu236, Met237, Asn239, Trp275 |
| **HG185** | A21C K50Q Q90F A125T W275V | –201.9 | 87.5 | 1 | 4 | 26 | Cys21, Gln50, Met237 |
| **HG198** | K50Q M84C A125T F267M | –201.6 | 99.8 | 1 | 4 | 23 | Trp44, Gln50, Thr125, Gln207, Met237, Met267 |
| **HG259** | K50Q M84C Q90L A125T F267M | –200.5 | 99.8 | 1 | 4 | 23 | Trp44, Gln50, Thr125, Gln207, Met237, Met267 |
| **HG378** | A21C K50Q Q90L A125V W275V | –198.7 | 87.5 | 1 | 4 | 23 | Cys21, Trp44, Gln50, Val125, Leu236, Met237 |
| **HG493** | K50Q Q90F A125T W275T | –197.1 | 87.1 | 0 | 4 | 24 | Trp44, Gln50, Thr125, Leu236, Met237 |
| **HG603** | K50Q Q90F A125V W275V | –195.5 | 87.1 | 0 | 4 | 24 | Trp44, Gln50, Thr125, Leu236, Met237 |
| **HG630** | K50Q M84C A125T F267M W275V | –194.9 | 99.8 | 1 | 4 | 22 | Trp44, Asn47, Gln50, Thr125, Gln207, Met237, Met267 |
| **HG649** | K50Q M84C Q90F A125T F267M W275V | –194.6 | 99.8 | 1 | 4 | 23 | Trp44, Gln50, Thr125, Gln207, Met237, Met267 |

[a] Solvent-accessible surface area of the transition state in the design model.
[b] Preorganized residues are those predicted to adopt the same rotamer in the presence and absence of transition state.

**Supplementary Table 3.** Angles defining the hydrogen bond between D127 and the transition state or transition-state analogue for various Kemp eliminase structures

| Template | PDB ID | Angle (°) | | | Deviation from ideal (°) | | |
|---|---|---|---|---|---|---|---|
| | | $\psi$ [a] | $\theta$ [a] | $\chi$ [a] | $\psi$ | $\theta$ | $\chi$ |
| **Crystal structures**[b] | | | | | | | |
| **HG3** | 5RGA | 134 ± 1 | 173 ± 2 | 182 ± 3 | 22 | 14 | 4.5 |
| **HG3.7** | 5RGC | 125 ± 1 | 167 ± 1 | 174 ± 2 | 13 | 8 | 3.5 |
| **HG185** | 8USL | 125 ± 3 | 163 ± 3 | 175 ± 9 | 13 | 4 | 2.5 |
| **HG198** | 8USJ | 124 ± 2 | 168 ± 1 | 186 ± 5 | 12 | 9 | 8.5 |
| **HG630** | 8USH | 125 ± 1 | 170 ± 1 | 173 ± 5 | 13 | 11 | 4.5 |
| **HG649** | 8USF | 124 ± 2 | 169 ± 2 | 181 ± 9 | 12 | 10 | 3.5 |
| **Ensemble members**[c] | | | | | | | |
| **HG3 bb05** | – | 113.6 | 164.0 | 175.2 | 1.3 | 4.6 | 2.3 |
| **KE70 bb06** | – | 116.2 | 159.3 | 173.0 | 3.9 | 0.1 | 4.5 |

[a] Average angles across all chains are reported (mean ± s.d.). Ideal values for $\psi$, $\theta$ and $\chi$ are 112.3, 159.4 and 177.5 degrees, respectively. Atoms defining $\psi$, $\theta$ and $\chi$ are shown on Supplementary Figure 2.
[b] Crystal structures contain the 6NT transition-state analogue.
[c] Ensemble members contain the transition state.

**Supplementary Table 4.** Crystallization conditions

| Enzyme | 6NT | Method [a] | Buffer | pH | Protein (mg mL$^{-1}$) | Precipitant #1 | Precipitant #2 |
|---|---|---|---|---|---|---|---|
| **HG185** | (–) | Crystallization | 100 mM sodium acetate | 4.6 | 12.8 | 1.6 M ammonium sulfate | – |
| | (+) | Soaking | 100 mM sodium acetate | 4.6 | 12.8 | 1.6 M ammonium sulfate | – |
| **HG198** | (–) | Soaking | 100 mM sodium acetate | 4.6 | 6.0 | 1.8 M ammonium sulfate | – |
| | (+) | Co-crystallization | 100 mM sodium acetate | 4.6 | 6.0 | 1.6 M ammonium sulfate | – |
| **HG630** | (–) | Crystallization | 100 mM sodium acetate | 4.8 | 10.6 | 1.4 M ammonium sulfate | – |
| | (+) | Soaking | 100 mM sodium acetate | 4.8 | 10.6 | 1.4 M ammonium sulfate | – |
| **HG649** | (–) | Crystallization | – | n.d. | 12.5 | 0.3 M ammonium sulfate | 30% PEG-4000 |
| | (+) | Soaking | – | n.d. | 12.5 | 0.3 M ammonium sulfate | 30% PEG-4000 |

[a] Crystals of ligand-bound HG185, HG630 and HG649 were obtained by soaking ligand-free crystals in 1 µL drops of mother liquor containing 5 mM 6NT and 5% DMSO in a single step. Crystals of ligand-free HG198 were obtained by sequentially soaking crystals obtained by co-crystallization with 6NT in three new drops of 1 µL mother liquor containing no ligand (30 min incubation per drop).

**Supplementary Table 5.** Crystallographic data and refinement statistics

| 6NT | HG185 (–) | HG185 (+) | HG198 (–) | HG198 (+) | HG630 (–) | HG630 (+) | HG649 (–) | HG649 (+) |
|---|---|---|---|---|---|---|---|---|
| **PDB ID** | 8USK | 8USL | 8USI | 8USJ | 8USG | 8USH | 8USE | 8USF |
| **Data collection** [a] | | | | | | | | |
| **Temperature (K)** | 280 | 280 | 280 | 280 | 280 | 280 | 280 | 280 |
| **Resolution (Å)** | 1.60 | 1.55 | 1.90 | 1.60 | 1.37 | 1.32 | 1.55 | 1.45 |
| **Space group** | $P2_1$ | $P2_1$ | $P2_1$ | P1 | $P2_1$ | $P2_1$ | $P2_1$ | $P2_1$ |
| ***Cell params.*** | | | | | | | | |
| **a b c (Å)** | 51.12<br>77.44<br>93.75 | 51.11<br>77.49<br>93.85 | 45.68<br>58.72<br>51.30 | 46.65<br>51.03<br>86.30 | 51.12<br>77.51<br>93.88 | 51.02<br>77.51<br>93.90 | 51.13<br>77.47<br>93.78 | 51.07<br>77.47<br>93.81 |
| **α β γ (°)** | 90.00<br>105.55<br>90.00 | 90.00<br>105.51<br>90.00 | 90.00<br>95.36<br>90.00 | 83.58<br>81.12<br>87.01 | 90.00<br>105.42<br>90.00 | 90.00<br>105.27<br>90.00 | 90.00<br>105.39<br>90.00 | 90.00<br>105.28<br>90.00 |
| **Chains per asymm. unit** | 2 | 2 | 1 | 3 | 2 | 2 | 2 | 2 |
| $R_{pim}$ | 0.071<br>(0.542) | 0.065<br>(0.470) | 0.169<br>(0.517) | 0.123<br>(0.547) | 0.062<br>(0.443) | 0.052<br>(0.585) | 0.070<br>(0.513) | 0.059<br>(0.552) |
| $CC_{1/2}$ | 0.992<br>(0.474) | 0.994<br>(0.531) | 0.955<br>(0.493) | 0.981<br>(0.466) | 0.990<br>(0.551) | 0.995<br>(0.477) | 0.991<br>(0.505) | 0.996<br>(0.452) |
| **I/σI** | 10.3<br>(1.3) | 11.1<br>(1.5) | 6.0<br>(1.7) | 7.7<br>(1.7) | 8.2<br>(1.0) | 9.4<br>(1.1) | 7.9<br>(1.4) | 8.7<br>(1.0) |
| **Completeness (%)** | 96.4<br>(94.1) | 99.7<br>(99.4 | 99.9<br>(99.6) | 94.3<br>(91.8) | 99.4<br>(94.4) | 99.7<br>(98.9) | 99.4<br>(98.6) | 99.7<br>(99.7) |
| **Multiplicity** | 3.4<br>(3.5) | 3.3<br>(3.3) | 3.4<br>(3.1) | 1.8<br>(1.8) | 3.2<br>(2.7) | 3.3<br>(3.2) | 3.3<br>(3.1) | 3.2<br>(3.1) |
| **Wilson B-factor ($Å^2$)** | 14.880 | 14.190 | 9.580 | 10.130 | 14.910 | 13.060 | 14.580 | 12.510 |
| **# unique reflections** | 89399<br>(4304) | 101929<br>(5042) | 21360<br>(1030) | 97130<br>(4660) | 147144<br>(6954) | 164672<br>(8177) | 101581<br>(4971) | 124369<br>(6172) |
| **Refinement** | | | | | | | | |
| **R work/free** | 0.1605/<br>0.1833 | 0.1610/<br>0.1831 | 0.1585/<br>0.1991 | 0.1540/<br>0.1900 | 0.1274/<br>0.1408 | 0.1190/<br>0.1476 | 0.1471/<br>0.1727 | 0.1373/<br>0.1567 |
| ***No. atoms*** | | | | | | | | |
| **Protein** | 5068 | 5116 | 2408 | 7489 | 5156 | 5166 | 5188 | 5260 |
| **Ligand** | 0 | 24 | 0 | 36 | 10 | 24 | 0 | 24 |
| **Water** | 543 | 362 | 294 | 820 | 557 | 557 | 743 | 748 |
| ***Averaged B-factors ($Å^2$)*** | | | | | | | | |
| **Protein** | 19.74 | 19.04 | 14.02 | 14.54 | 19.37 | 17.87 | 19.77 | 17.50 |
| **Ligand** | – | 32.92 | – | 13.70 | 81.47 | 17.95 | – | 17.02 |
| **Water** | 35.18 | 29.85 | 27.68 | 30.53 | 36.59 | 39.27 | 37.66 | 35.87 |
| ***RMSD*** | | | | | | | | |
| **bond lengths (Å)** | 0.005 | 0.007 | 0.003 | 0.004 | 0.007 | 0.007 | 0.005 | 0.007 |
| **bond angles (°)** | 0.834 | 0.848 | 0.607 | 0.724 | 1.003 | 0.886 | 0.811 | 0.885 |
| ***Molprobity statistics*** | | | | | | | | |
| **Ramachand. outliers (%)** | 0.00 | 0.00 | 0.00 | 0.00 | 0.00 | 0.00 | 0.00 | 0.00 |
| **Ramachand. allowed (%)** | 1.67 | 1.51 | 1.66 | 1.56 | 1.51 | 1.34 | 1.84 | 1.17 |
| **Ramachan. favored (%)** | 98.33 | 98.49 | 98.34 | 98.44 | 98.49 | 98.66 | 98.16 | 98.83 |
| **Rotamer outliers (%)** | 0.00 | 0.00 | 0.00 | 0.00 | 0.00 | 0.00 | 0.00 | 0.00 |
| **MolProbity clashscore** | 4.15 | 1.67 | 1.67 | 1.73 | 2.52 | 2.32 | 3.57 | 3.03 |

[a] Highest resolution shell is shown in parentheses.

**Supplementary Table 6.** Structural comparison of bb05 and HG3 crystal structure with ensembles of HG-series variants

| | Median pairwise backbone RMSD from reference structure (Å)[a] | | | ΔRMSD from HG3 (Å) | | Z-score between distributions | |
|---|---|---|---|---|---|---|---|
| **Ensemble[b]** | **Self[c]** | **bb05[d]** | **HG3 xtal[e]** | **bb05[d]** | **HG3 xtal[e]** | **bb05[d]** | **HG3 xtal[e]** |
| **HG3** | 0.22 | 0.25 | 0.17 | 0 | 0 | – | – |
| **HG185** | 0.23 | 0.27 | 0.21 | +0.02 | +0.04 | –5.7 | –7.1 |
| **HG198** | 0.17 | 0.27 | 0.23 | +0.02 | +0.06 | –5.4 | –10.6 |
| **HG630** | 0.12 | 0.26 | 0.22 | +0.01 | +0.05 | –1.1 | –9.2 |
| **HG649** | 0.09 | 0.23 | 0.19 | –0.02 | +0.02 | +2.0 | –4.9 |

[a] Median pairwise backbone RMSDs were calculated for designed active-site residues listed on Supplementary Table 7.
[b] Ensembles were generated by ensemble refinement of 6NT-bound crystal structures listed on Supplementary Table 1.
[c] Median pairwise backbone RMSD between every pair of backbones from the same ensemble.
[d] HG3 (+) 6NT ensemble member used to design HG185, HG198, HG630 and HG649.
[e] HG3 crystal structure bound with 6NT (PDB ID: 5RGA, Chain A).

**Supplementary Table 7.** Amino-acid positions optimized during computational design of HG3 variants

| Design objective | Designed residues [a] | Allowed amino acids |
|---|---|---|
| **Catalytic base** | D127 | WT |
| **H-bond donor** | A21 W44 K50 M84 W87 Q90 A125 Y170 M172 L236 M237 T265 F267 | N, Q, S, T |
| **Transition-state packing** | A21 W44 M84 W87 Q90 A125 Y170 M172 L236 M237 T265 F267 | A, C, F, I, L, M, V, T, WT [b] |
| **Active-site bottleneck [c]** | W275 | A, S, V, T, WT [b] |
| **Floated positions [d]** | V16 Y17 M42 E46 N47 L79 G130 Q207 H209 S234 D239 R276 | WT [b] |

[a] Numbering based on crystal structure of HG3 (PDB ID: 5RGA).
[b] WT indicates the amino acid found at that position in the HG3 sequence (e.g., N, Q, and Y at positions 47, 90, and 170, respectively)
[c] The bulky W275 residue was mutated to small amino acids to widen the active-site entrance bottleneck.
[d] Amino acids at these positions were allowed to change conformation but not identity.

**Supplementary Table 8.** Design filtering – KE70 series

| Design | Mutations from KE70 | Energy (kcal mol$^{-1}$) | SASA (Å$^2$) [a] | # Cys | # Met | # Preorganized residues [b] | Residues not preorganized [c] |
|---|---|---|---|---|---|---|---|
| **KE701** | H16D S20a D45Q Y48F W72C G101S S138Q H166N V168A A204V A240a | –29.4 | 101.0 | 1 | 0 | 12 | Phe48, Trp72, Gln138 |
| **KE702** | H16D S20a D45Q Y48F R70K W72Y G101M S138Q H166N V168A I202V A204M A240a | –17.9 | 41.4 | 0 | 2 | 12 | Asp16, Gln138, Met204 |
| **KE703** | H16D S20a D45N Y48F W72V G101M S138Q H166N V168A A204V A240a | –17.8 | 101.6 | 0 | 1 | 12 | Phe48, Trp72, Gln138 |

[a] Solvent-accessible surface area of the transition state in the design model.
[b] Preorganized residues are those predicted to adopt the same rotamer in the presence and absence of transition state.
[c] Numbering based on sequence shown on Supplementary Table 13 and Supplementary Figure 8.

**Supplementary Table 9.** Amino-acid positions optimized during computational design of KE70 variants

| Design objective | Designed residues [a] | Allowed amino acids |
|---|---|---|
| **Catalytic base** | H17 [b] | D |
| **H-bond donor** | S138 | Q |
| **Pi-stacking** | Y48 | F |
| **Removal of original catalytic dyad** [c] | D45 | N, Q |
| **Transition-state packing** [d] | A19 W72 G101 A103 I140 H166 V168 S170 I202 A204 | A, F, I, L, M, N, V, Y |
| **Surface residue** [e] | R70 | K, R |

[a] Numbering based on crystal structure of KE70 (PDB ID: 3NPV). For comparison with sequences of designed variants, see Supplementary Figure 8.
[b] Position 17 in the 3NPV structure correspond to position 16 in the sequence of our designed variants. See sequence alignment on Supplementary Figure 8.
[c] KE70 contains the H17/D45 catalytic dyad. The D45N and D45Q mutations were introduced to make this residue compatible with the newly designed catalytic base (D16).
[d] C and S were also allowed at positions W72 and G101 instead of A.
[e] This surface residue is located next to D45.

**Supplementary Table 10.** Geometric definitions for generation of transition-state poses off the side chain of catalytic base

| **Residue** | **Type** | **Atom 1** [a] | **Atom 2** [a] | **Atom 3** [a] | **Atom 4** [a] | **Values** [b] |
|---|---|---|---|---|---|---|
| **Asp** | Distance | OD1 or OD2 | **H3** | | | 1.0, 1.2, 1.5 |
| | Angle | CG | OD1 or OD2 | **H3** | | 112, 117, 122 |
| | Angle | OD1 or OD2 | **H3** | **C3** | | 159, 164, 169, 174, 179 |
| | Torsion | CB | CG | OD1 or OD2 | **H3** | 0, 5, 10, 170, 175, 180 |
| | Torsion | CG | OD1 or OD2 | **H3** | **C3** | 170, 175, 180, 185, 190 |
| | Torsion | OD1 or OD2 | **H3** | **C3** | **N2** | 0, 5, 170, 175, 180 |

[a] Atoms in bold are from the transition state. All other atoms are from the catalytic residue.
[b] Distance measurements given in Å, all others are in degrees.

**Supplementary Table 11.** Geometric constraints used to define catalytic contacts during computational design

| Contact | Residue | Type | Atom 1 [a] | Atom 2 [a] | Atom 3 [a] | Atom 4 [a] | Min [b] | Max [b] |
|---|---|---|---|---|---|---|---|---|
| **Catalytic base** | Asp | Distance | OD1 or OD2 | **H3** | | | 1.0 (1.0) | 1.6 (1.6) |
| | | Angle | CG | OD1 or OD2 | **H3** | | 109 (109) | 131 (131) |
| | | Angle | OD1 or OD2 | **H3** | **C3** | | 159 (159) | 180 (180) |
| | | Torsion | CB | CG | OD1 or OD2 | **H3** | –21 (–21) | 21 (21) |
| **H-bond donor** | Gln | Distance | 1HE2 or 2HE2 | **O1** | | | 1.2 (1.2) | 2.3 (2.3) |
| | | Angle | NE2 | 1HE2 or 2HE2 | **O1** | | 145 (145) | 157 (157) |
| | | Angle | 1HE2 or 2HE2 | **O1** | **N2** | | 120 (120) | 140 (140) |
| | | Torsion | 1HE2 or 2HE2 | O1 | N2 | C3 | 160 (160) | 200 (200) |
| | Asn | Distance | 1HD2 or 2HD2 | **O1** | | | 1.2 (1.2) | 2.3 (2.3) |
| | | Angle | ND2 | 1HD2 or 2HD2 | **O1** | | 145 (145) | 157 (157) |
| | | Angle | 1HD2 or 2HD2 | **O1** | **N2** | | 120 (120) | 140 (140) |
| | | Torsion | 1HD2 or 2HD2 | O1 | N2 | C3 | 160 (160) | 200 (200) |
| | Thr | Distance | HG1 | **O1** | | | 1.2 (1.2) | 2.3 (2.3) |
| | | Angle | OG1 | HG1 | **O1** | | 145 (145) | 157 (157) |
| | | Angle | HG1 | **O1** | **N2** | | 120 (120) | 140 (140) |
| | | Torsion | HG1 | O1 | N2 | C3 | 160 (160) | 200 (200) |
| | Ser | Distance | HG | **O1** | | | 1.2 (1.2) | 2.3 (2.3) |
| | | Angle | OG | HG | **O1** | | 145 (145) | 157 (157) |
| | | Angle | HG | **O1** | **N2** | | 120 (120) | 140 (140) |
| | | Torsion | HG | O1 | N2 | C3 | 160 (160) | 200 (200) |
| **Pi-stacking** [c] | Phe | Distance between aromatic planes | CE1 CD2 | **C4 C9** | | | 3.5 (3.5) | 4 (4) |
| | | Angle between aromatic planes | CG CE1 CR2 | **C5 C8 N2** | | | 0 (120) | 40 (180) |

[a] Atoms in bold are from the transition state. All other atoms are from the catalytic residues.
[b] Distance measurements given in Å, all others in degrees. Values are for theozyme placement, while those in parentheses are for active-site repacking.
[c] This catalytic contact was used for KE70 designs only.

**Supplementary Table 12.** Amino-acid sequences of Kemp eliminases from HG series

| Enzyme | # Mutations from HG3 | Sequences |
|---|---|---|
| **HG44** | 4 | MAEAAQSVDQLIKARGKVYFGVCTDQNRLTTGKNAAIIQADFGMVWPENSMQWDATEPSQGNFNFAGADYLVNWAQQN<br>GKLIGGGMLVWHSFLPSWVSSITDKNTLTNVMKNHITTLMTRYKGKIRVWDVVGEAFNEDGSLRQTVFLNVIGEDYIP<br>IAFQTARAADPNAKLYIMDYNLDSASYPKTQAIVNRVKQWRAAGVPIDGIGSQTHLSAGQGAGVLQALPLLASAGTPE<br>VSILMLDVAGASPTDYVNVVNACLNVQSCVGITVFGVADPDSWRASTTPLLFDGNFNPKPAYNAIVQDLQQGSIEGRG<br>HHHHHH |
| **HG57** | 4 | MAEAAQSVDQLIKARGKVYFGVCTDQNRLTTGKNAAIIQADFGMVWPENSMQWDATEPSQGNFNFAGADYLVNWAQQN<br>GKLIGGGMLVWHSLLPSWVSSITDKNTLTNVMKNHITTLMTRYKGKIRVWDVVGEAFNEDGSLRQTVFLNVIGEDYIP<br>IAFQTARAADPNAKLYIMDYNLDSASYPKTQAIVNRVKQWRAAGVPIDGIGSQTHLSAGQGAGVLQALPLLASAGTPE<br>VSILMLDVAGASPTDYVNVVNACLNVQSCVGITVFGVADPDSWRASTTPLLFDGNFNPKPAYNAIVQDLQQGSIEGRG<br>HHHHHH |
| **HG185** | 5 | MAEAAQSVDQLIKARGKVYFGVCTDQNRLTTGKNAAIIQADFGMVWPENSMQWDATEPSQGNFNFAGADYLVNWAQQN<br>GKLIGGGMLVWHSFLPSWVSSITDKNTLTNVMKNHITTLMTRYKGKIRTWDVVGEAFNEDGSLRQTVFLNVIGEDYIP<br>IAFQTARAADPNAKLYIMDYNLDSASYPKTQAIVNRVKQWRAAGVPIDGIGSQTHLSAGQGAGVLQALPLLASAGTPE<br>VSILMLDVAGASPTDYVNVVNACLNVQSCVGITVFGVADPDSVRASTTPLLFDGNFNPKPAYNAIVQDLQQGSIEGRG<br>HHHHHH |
| **HG198** | 4 | MAEAAQSVDQLIKARGKVYFGVATDQNRLTTGKNAAIIQADFGMVWPENSMQWDATEPSQGNFNFAGADYLVNWAQQN<br>GKLIGGGCLVWHSQLPSWVSSITDKNTLTNVMKNHITTLMTRYKGKIRTWDVVGEAFNEDGSLRQTVFLNVIGEDYIP<br>IAFQTARAADPNAKLYIMDYNLDSASYPKTQAIVNRVKQWRAAGVPIDGIGSQTHLSAGQGAGVLQALPLLASAGTPE<br>VSILMLDVAGASPTDYVNVVNACLNVQSCVGITVMGVADPDSWRASTTPLLFDGNFNPKPAYNAIVQDLQQGSIEGRG<br>HHHHHH |
| **HG259** | 5 | MAEAAQSVDQLIKARGKVYFGVATDQNRLTTGKNAAIIQADFGMVWPENSMQWDATEPSQGNFNFAGADYLVNWAQQN<br>GKLIGGGCLVWHSLLPSWVSSITDKNTLTNVMKNHITTLMTRYKGKIRTWDVVGEAFNEDGSLRQTVFLNVIGEDYIP<br>IAFQTARAADPNAKLYIMDYNLDSASYPKTQAIVNRVKQWRAAGVPIDGIGSQTHLSAGQGAGVLQALPLLASAGTPE<br>VSILMLDVAGASPTDYVNVVNACLNVQSCVGITVMGVADPDSWRASTTPLLFDGNFNPKPAYNAIVQDLQQGSIEGRG<br>HHHHHH |
| **HG378** | 5 | MAEAAQSVDQLIKARGKVYFGVCTDQNRLTTGKNAAIIQADFGMVWPENSMQWDATEPSQGNFNFAGADYLVNWAQQN<br>GKLIGGGMLVWHSLLPSWVSSITDKNTLTNVMKNHITTLMTRYKGKIRVWDVVGEAFNEDGSLRQTVFLNVIGEDYIP<br>IAFQTARAADPNAKLYIMDYNLDSASYPKTQAIVNRVKQWRAAGVPIDGIGSQTHLSAGQGAGVLQALPLLASAGTPE<br>VSILMLDVAGASPTDYVNVVNACLNVQSCVGITVFGVADPDSVRASTTPLLFDGNFNPKPAYNAIVQDLQQGSIEGRG<br>HHHHHH |
| **HG493** | 4 | MAEAAQSVDQLIKARGKVYFGVATDQNRLTTGKNAAIIQADFGMVWPENSMQWDATEPSQGNFNFAGADYLVNWAQQN<br>GKLIGGGMLVWHSFLPSWVSSITDKNTLTNVMKNHITTLMTRYKGKIRTWDVVGEAFNEDGSLRQTVFLNVIGEDYIP<br>IAFQTARAADPNAKLYIMDYNLDSASYPKTQAIVNRVKQWRAAGVPIDGIGSQTHLSAGQGAGVLQALPLLASAGTPE<br>VSILMLDVAGASPTDYVNVVNACLNVQSCVGITVFGVADPDSTRASTTPLLFDGNFNPKPAYNAIVQDLQQGSIEGRG<br>HHHHHH |
| **HG603** | 4 | MAEAAQSVDQLIKARGKVYFGVATDQNRLTTGKNAAIIQADFGMVWPENSMQWDATEPSQGNFNFAGADYLVNWAQQN<br>GKLIGGGMLVWHSFLPSWVSSITDKNTLTNVMKNHITTLMTRYKGKIRVWDVVGEAFNEDGSLRQTVFLNVIGEDYIP<br>IAFQTARAADPNAKLYIMDYNLDSASYPKTQAIVNRVKQWRAAGVPIDGIGSQTHLSAGQGAGVLQALPLLASAGTPE<br>VSILMLDVAGASPTDYVNVVNACLNVQSCVGITVFGVADPDSVRASTTPLLFDGNFNPKPAYNAIVQDLQQGSIEGRG<br>HHHHHH |
| **HG630** | 5 | MAEAAQSVDQLIKARGKVYFGVATDQNRLTTGKNAAIIQADFGMVWPENSMQWDATEPSQGNFNFAGADYLVNWAQQN<br>GKLIGGGCLVWHSQLPSWVSSITDKNTLTNVMKNHITTLMTRYKGKIRTWDVVGEAFNEDGSLRQTVFLNVIGEDYIP<br>IAFQTARAADPNAKLYIMDYNLDSASYPKTQAIVNRVKQWRAAGVPIDGIGSQTHLSAGQGAGVLQALPLLASAGTPE<br>VSILMLDVAGASPTDYVNVVNACLNVQSCVGITVMGVADPDSVRASTTPLLFDGNFNPKPAYNAIVQDLQQGSIEGRG<br>HHHHHH |
| **HG649** | 6 | MAEAAQSVDQLIKARGKVYFGVATDQNRLTTGKNAAIIQADFGMVWPENSMQWDATEPSQGNFNFAGADYLVNWAQQN<br>GKLIGGGCLVWHSFLPSWVSSITDKNTLTNVMKNHITTLMTRYKGKIRTWDVVGEAFNEDGSLRQTVFLNVIGEDYIP<br>IAFQTARAADPNAKLYIMDYNLDSASYPKTQAIVNRVKQWRAAGVPIDGIGSQTHLSAGQGAGVLQALPLLASAGTPE<br>VSILMLDVAGASPTDYVNVVNACLNVQSCVGITVMGVADPDSVRASTTPLLFDGNFNPKPAYNAIVQDLQQGSIEGRG<br>HHHHHH |

**Supplementary Table 13.** Amino-acid sequences of Kemp eliminases from KE70 series

| Enzyme | # Mutations from KE70 | Sequence |
|---|---|---|
| **KE70_S20a A240a** | 2 | MTDLKASSLRALKLMHLATSANDDDTDEKVIALCHQAKTPVGTTDAIYIYPRFIPIARKTLKEQGTPEIRI<br>WTSTNFPHGNDDIDIALAETRAAIAYGADGVAVVFPYRALMAGNEQVGFDLVKACKEACAAANVLLSVIIE<br>TGELKDEALIRKASEISIKAGADHIVTSTGKVAVGATPESARIMMEVIRDMGVEKTVGFIPAGGVRTAEDA<br>QKYLAIADELFGADWADARHYAFGASASLLASLLKALGHGDGKSASSYGSLEHHHHHH |
| **KE701** | 11 | MTDLKASSLRALKLMDLATSANDDDTDEKVIALCHQAKTPVGTTQAIFIYPRFIPIARKTLKEQGTPEIRI<br>CTSTNFPHGNDDIDIALAETRAAIAYGADSVAVVFPYRALMAGNEQVGFDLVKACKEACAAANVLLQVIIE<br>TGELKDEALIRKASEISIKAGADNIATSTGKVAVGATPESARIMMEVIRDMGVEKTVGFIPVGGVRTAEDA<br>QKYLAIADELFGADWADARHYAFGASASLLASLLKALGHGDGKSASSYGSLEHHHHHH |
| **KE702** | 13 | MTDLKASSLRALKLMDLATSANDDDTDEKVIALCHQAKTPVGTTNAIFIYPRFIPIARKTLKEQGTPEIKI<br>YTSTNFPHGNDDIDIALAETRAAIAYGADMVAVVFPYRALMAGNEQVGFDLVKACKEACAAANVLLQVIIE<br>TGELKDEALIRKASEISIKAGADNIATSTGKVAVGATPESARIMMEVIRDMGVEKTVGFVPMGGVRTAEDA<br>QKYLAIADELFGADWADARHYAFGASASLLASLLKALGHGDGKSASSYGSLEHHHHHH |
| **KE703** | 11 | MTDLKASSLRALKLMDLATSANDDDTDEKVIALCHQAKTPVGTTNAIFIYPRFIPIARKTLKEQGTPEIRI<br>VTSTNFPHGNDDIDIALAETRAAIAYGADMVAVVFPYRALMAGNEQVGFDLVKACKEACAAANVLLQVIIE<br>TGELKDEALIRKASEISIKAGADNIATSTGKVAVGATPESARIMMEVIRDMGVEKTVGFIPVGGVRTAEDA<br>QKYLAIADELFGADWADARHYAFGASASLLASLLKALGHGDGKSASSYGSLEHHHHHH |
| **KE703b** | 9 | MTDLKASSLRALKLMHLATSANDDDTDEKVIALCHQAKTPVGTTDAIFIYPRFIPIARKTLKEQGTPEIRI<br>VTSTNFPHGNDDIDIALAETRAAIAYGADMVAVVFPYRALMAGNEQVGFDLVKACKEACAAANVLLQVIIE<br>TGELKDEALIRKASEISIKAGADNIATSTGKVAVGATPESARIMMEVIRDMGVEKTVGFIPVGGVRTAEDA<br>QKYLAIADELFGADWADARHYAFGASASLLASLLKALGHGDGKSASSYGSLEHHHHHH |

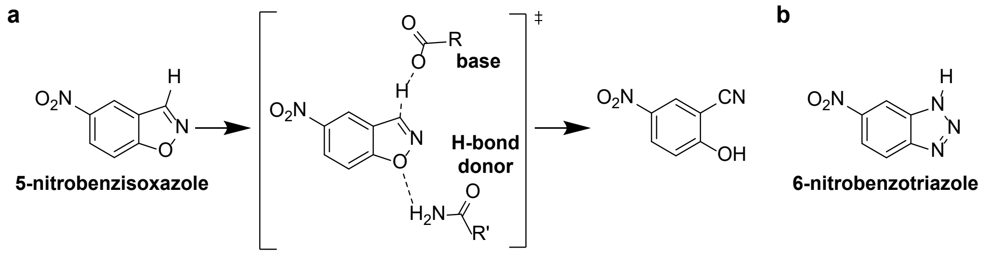


**Supplementary Figure 1. The Kemp elimination reaction.** (a) HG-series Kemp eliminases catalyze the Kemp elimination using a catalytic motif consisting of a base (Asp) that deprotonates the 5-nitrobenzisoxazole substrate, and an H-bond donor (Gln) that stabilizes negative charge buildup on the phenolic oxygen at the transition state (‡). This reaction yields the 4-nitro-2-cyanophenol product. (b) Structure of the 6NT transition-state analogue.

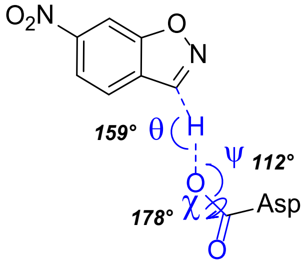


**Supplementary Figure 2. Ideal geometry for catalytic base contact.** Atoms defining various angles describing the hydrogen-bond interaction between the transition state and designed Asp catalytic base (θ, ψ, χ) are shown in blue. Values in italics are optimal angles calculated for hydrogen bonding interactions between acetamide dimers.

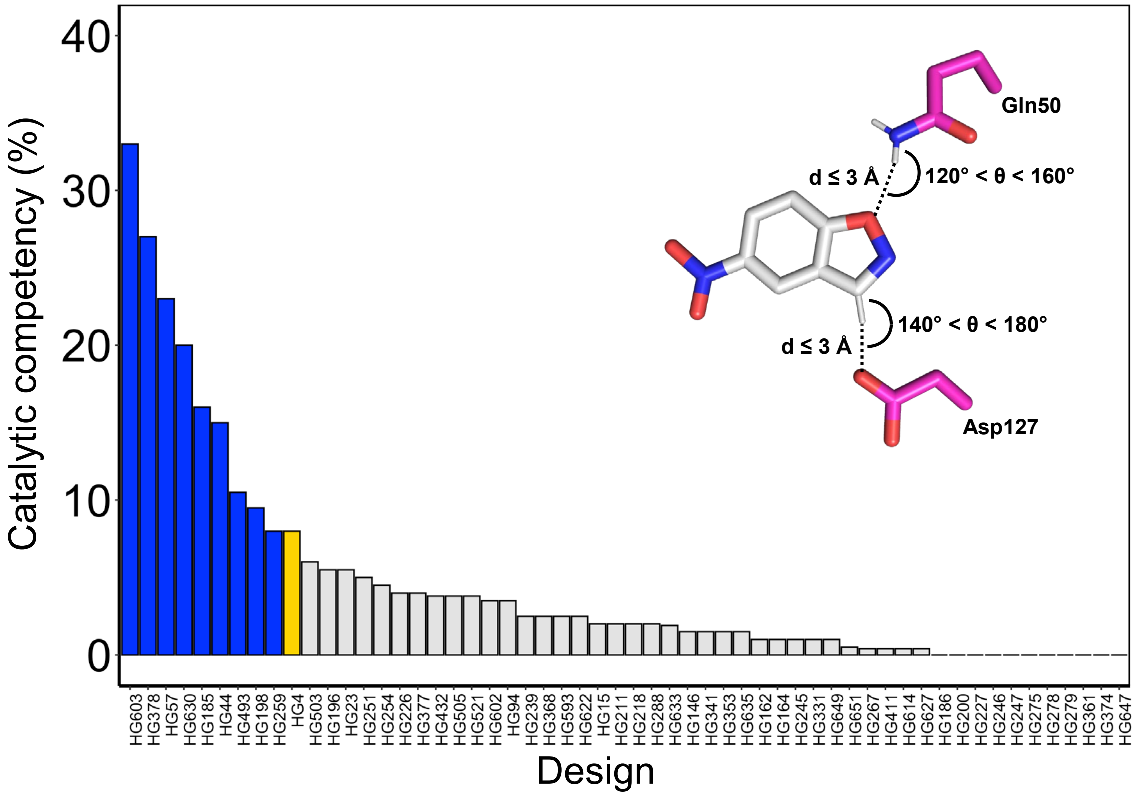


**Supplementary Figure 3. Design filtering using molecular dynamics.** Following molecular dynamics, angles (θ) and distances (d) defining catalytic contacts between the 5-nitrobenzisoxazole substrate and the designed catalytic residues (Asp127 and Gln50) were calculated on each snapshot. Snapshots whose θ and d values fall within the allowed ranges were considered to be catalytically competent. Catalytic competency is the percentage of snapshots in the trajectory that are catalytically competent. Only 9 variants (blue) have a catalytic competency that is greater than that of our positive control HG4 (yellow, PDB ID: 5RGF), an HG3 variant that is approximately 700-fold more catalytically efficient than HG3[14].

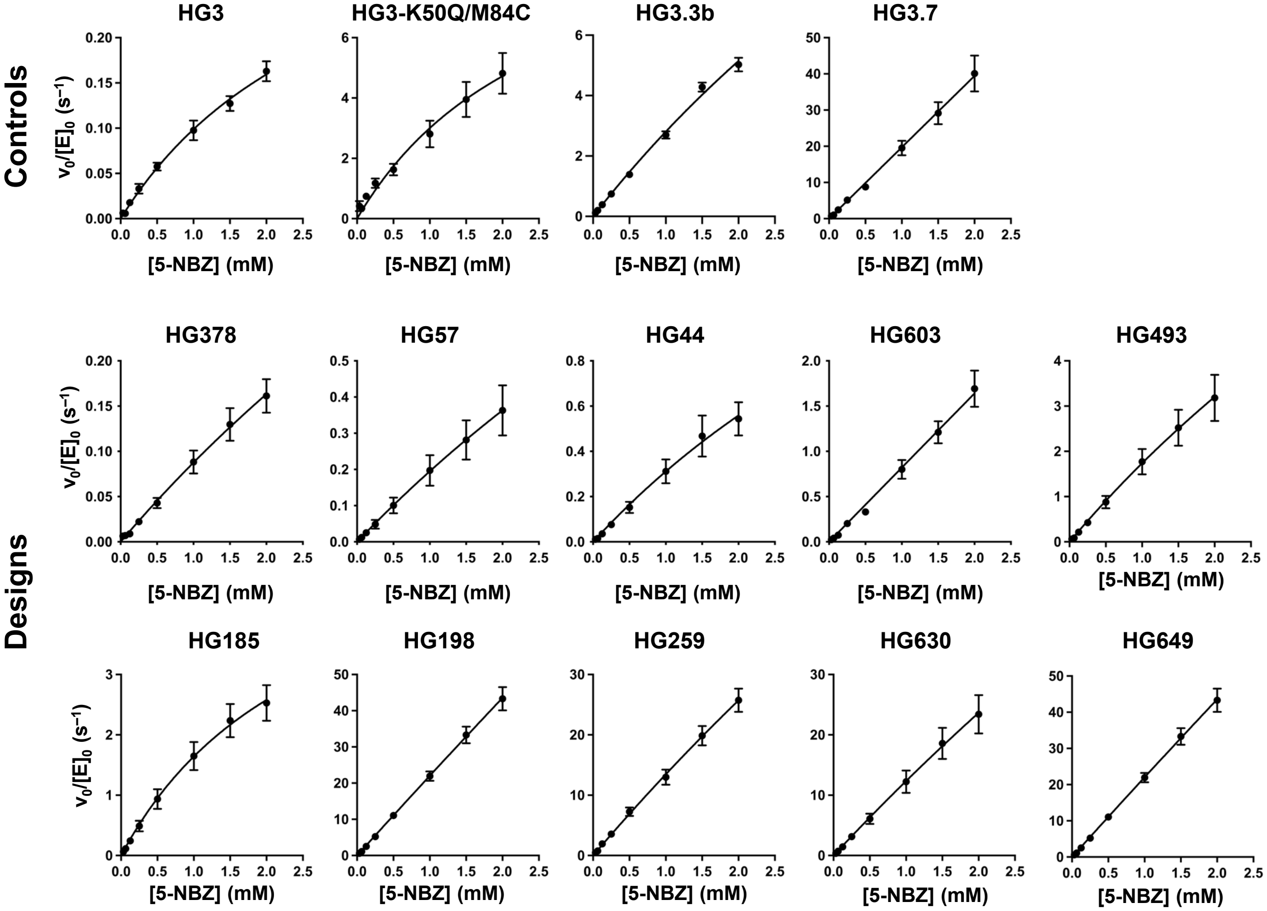


**Supplementary Figure 4. Steady-state kinetics of HG-series Kemp eliminases.** Michaelis–Menten plots of normalized initial rates as a function of 5-nitrobenzisoxazole (5-NBZ) concentrations are shown. Data represent the average of six or nine individual replicate measurements from two or three independent protein batches, with error bars indicating the SEM. Saturation was not achieved for any enzyme at the substrate's solubility limit (2 mM). Therefore, only $k_{cat}/K_M$ values are reported on Table 1, and these values were calculated using linear regression of rates measured at the three or four lowest substrate concentrations.

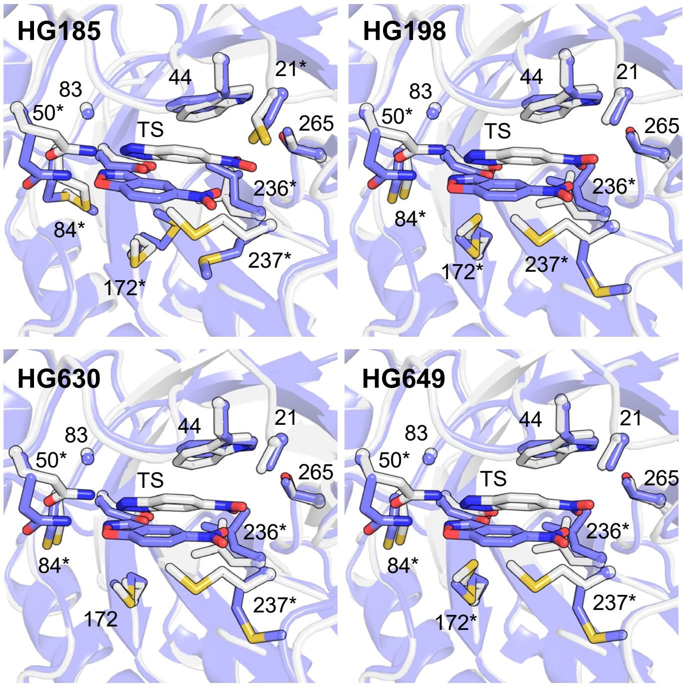


**Supplementary Figure 5. Design on HG3 average crystal structure does not accurately predict structures of designed enzymes.** Crystal structures of HG-series Kemp eliminases with bound 6NT (white) are overlaid on the corresponding design models (blue) obtained using the average crystal structure (PDB ID: 5RGA) as template. In all cases, the transition state (TS) and transition-state analogue are shown at the center of the image. Side chains of all residues forming the binding pocket are shown with the exception of P45, which was omitted for clarity. The sphere shows the alpha carbon of G83. Asterisks indicate residues that adopt side-chain rotamers varying by >20 degrees around one or more side-chain dihedrals between the design model and crystal structure. In all design models generated from the average crystal structure, the catalytic hydrogen-bond donor Q50 is predicted to adopt an alternate rotamer, which causes an erroneous prediction of the transition-state binding pose in the active site.

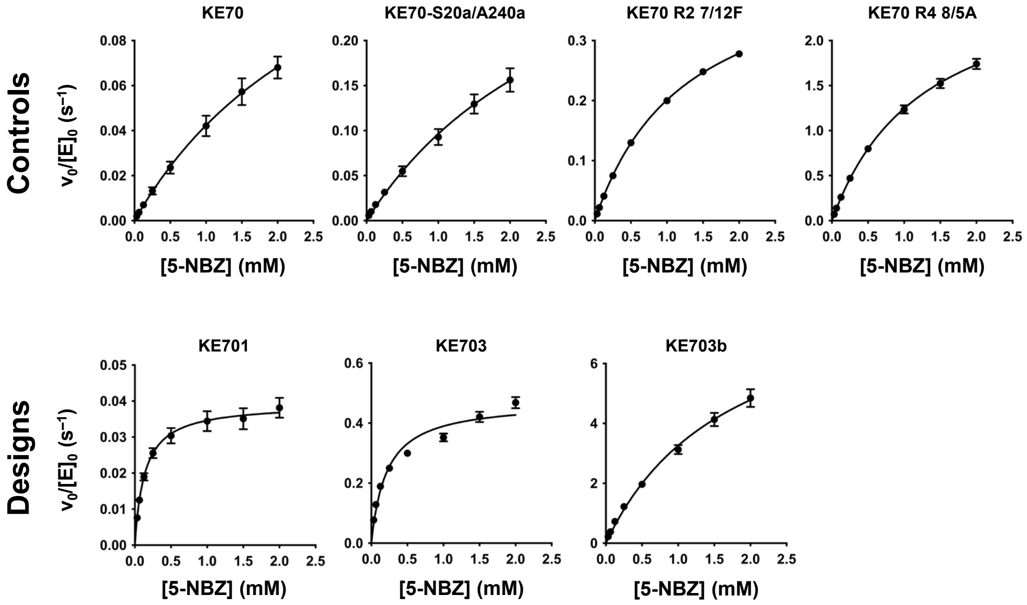


**Supplementary Figure 6. Steady-state kinetics of KE70-series Kemp eliminases.** Michaelis–Menten plots of normalized initial rates as a function of 5-nitrobenzisoxazole (5-NBZ) concentrations are shown. Data represent the average of six to twelve individual replicate measurements from two to four independent protein batches, with error bars indicating the SEM. Saturation was not achieved for some enzymes at the substrate's solubility limit (2 mM). In these cases, only $k_{cat}/K_M$ values are reported on Table 1, and these values were calculated using linear regression of rates measured at the three or four lowest substrate concentrations.

```
          1 ........10........20........30........40........50........60........70
HG3       1 MAEAAQSVDQLIKARGKVYFGVATDQNRLTTGKNAAIIQADFGMVWPENSMKWDATEPSQGNFNFAGADY
HG44      1 MAEAAQSVDQLIKARGKVYFGVCTDQNRLTTGKNAAIIQADFGMVWPENSMQWDATEPSQGNFNFAGADY
HG57      1 MAEAAQSVDQLIKARGKVYFGVCTDQNRLTTGKNAAIIQADFGMVWPENSMQWDATEPSQGNFNFAGADY
HG185     1 MAEAAQSVDQLIKARGKVYFGVCTDQNRLTTGKNAAIIQADFGMVWPENSMQWDATEPSQGNFNFAGADY
HG198     1 MAEAAQSVDQLIKARGKVYFGVATDQNRLTTGKNAAIIQADFGMVWPENSMQWDATEPSQGNFNFAGADY
HG259     1 MAEAAQSVDQLIKARGKVYFGVATDQNRLTTGKNAAIIQADFGMVWPENSMQWDATEPSQGNFNFAGADY
HG378     1 MAEAAQSVDQLIKARGKVYFGVCTDQNRLTTGKNAAIIQADFGMVWPENSMQWDATEPSQGNFNFAGADY
HG493     1 MAEAAQSVDQLIKARGKVYFGVATDQNRLTTGKNAAIIQADFGMVWPENSMQWDATEPSQGNFNFAGADY
HG603     1 MAEAAQSVDQLIKARGKVYFGVATDQNRLTTGKNAAIIQADFGMVWPENSMQWDATEPSQGNFNFAGADY
HG630     1 MAEAAQSVDQLIKARGKVYFGVATDQNRLTTGKNAAIIQADFGMVWPENSMQWDATEPSQGNFNFAGADY
HG649     1 MAEAAQSVDQLIKARGKVYFGVATDQNRLTTGKNAAIIQADFGMVWPENSMQWDATEPSQGNFNFAGADY

         71 ........80........90.......100.......110.......120.......130.......140
HG3      71 LVNWAQQNGKLIGGGMLVWHSQLPSWVSSITDKNTLTNVMKNHITTLMTRYKGKIRAWDVVGEAFNEDGS
HG44     71 LVNWAQQNGKLIGGGMLVWHSFLPSWVSSITDKNTLTNVMKNHITTLMTRYKGKIRVWDVVGEAFNEDGS
HG57     71 LVNWAQQNGKLIGGGMLVWHSLLPSWVSSITDKNTLTNVMKNHITTLMTRYKGKIRVWDVVGEAFNEDGS
HG185    71 LVNWAQQNGKLIGGGMLVWHSFLPSWVSSITDKNTLTNVMKNHITTLMTRYKGKIRTWDVVGEAFNEDGS
HG198    71 LVNWAQQNGKLIGGGCLVWHSQLPSWVSSITDKNTLTNVMKNHITTLMTRYKGKIRTWDVVGEAFNEDGS
HG259    71 LVNWAQQNGKLIGGGCLVWHSLLPSWVSSITDKNTLTNVMKNHITTLMTRYKGKIRTWDVVGEAFNEDGS
HG378    71 LVNWAQQNGKLIGGGMLVWHSLLPSWVSSITDKNTLTNVMKNHITTLMTRYKGKIRVWDVVGEAFNEDGS
HG493    71 LVNWAQQNGKLIGGGMLVWHSFLPSWVSSITDKNTLTNVMKNHITTLMTRYKGKIRTWDVVGEAFNEDGS
HG603    71 LVNWAQQNGKLIGGGMLVWHSFLPSWVSSITDKNTLTNVMKNHITTLMTRYKGKIRVWDVVGEAFNEDGS
HG630    71 LVNWAQQNGKLIGGGCLVWHSQLPSWVSSITDKNTLTNVMKNHITTLMTRYKGKIRTWDVVGEAFNEDGS
HG649    71 LVNWAQQNGKLIGGGCLVWHSFLPSWVSSITDKNTLTNVMKNHITTLMTRYKGKIRTWDVVGEAFNEDGS

        141 .......150.......160.......170.......180.......190.......200.......210
HG3     141 LRQTVFLNVIGEDYIPIAFQTARAADPNAKLYIMDYNLDSASYPKTQAIVNRVKQWRAAGVPIDGIGSQT
HG44    141 LRQTVFLNVIGEDYIPIAFQTARAADPNAKLYIMDYNLDSASYPKTQAIVNRVKQWRAAGVPIDGIGSQT
HG57    141 LRQTVFLNVIGEDYIPIAFQTARAADPNAKLYIMDYNLDSASYPKTQAIVNRVKQWRAAGVPIDGIGSQT
HG185   141 LRQTVFLNVIGEDYIPIAFQTARAADPNAKLYIMDYNLDSASYPKTQAIVNRVKQWRAAGVPIDGIGSQT
HG198   141 LRQTVFLNVIGEDYIPIAFQTARAADPNAKLYIMDYNLDSASYPKTQAIVNRVKQWRAAGVPIDGIGSQT
HG259   141 LRQTVFLNVIGEDYIPIAFQTARAADPNAKLYIMDYNLDSASYPKTQAIVNRVKQWRAAGVPIDGIGSQT
HG378   141 LRQTVFLNVIGEDYIPIAFQTARAADPNAKLYIMDYNLDSASYPKTQAIVNRVKQWRAAGVPIDGIGSQT
HG493   141 LRQTVFLNVIGEDYIPIAFQTARAADPNAKLYIMDYNLDSASYPKTQAIVNRVKQWRAAGVPIDGIGSQT
HG603   141 LRQTVFLNVIGEDYIPIAFQTARAADPNAKLYIMDYNLDSASYPKTQAIVNRVKQWRAAGVPIDGIGSQT
HG630   141 LRQTVFLNVIGEDYIPIAFQTARAADPNAKLYIMDYNLDSASYPKTQAIVNRVKQWRAAGVPIDGIGSQT
HG649   141 LRQTVFLNVIGEDYIPIAFQTARAADPNAKLYIMDYNLDSASYPKTQAIVNRVKQWRAAGVPIDGIGSQT

        211 .......220.......230.......240.......250.......260.......270.......280
HG3     211 HLSAGQGAGVLQALPLLASAGTPEVSILMLDVAGASPTDYVNVVNACLNVQSCVGITVFGVADPDSWRAS
HG44    211 HLSAGQGAGVLQALPLLASAGTPEVSILMLDVAGASPTDYVNVVNACLNVQSCVGITVFGVADPDSWRAS
HG57    211 HLSAGQGAGVLQALPLLASAGTPEVSILMLDVAGASPTDYVNVVNACLNVQSCVGITVFGVADPDSWRAS
HG185   211 HLSAGQGAGVLQALPLLASAGTPEVSILMLDVAGASPTDYVNVVNACLNVQSCVGITVFGVADPDSVRAS
HG198   211 HLSAGQGAGVLQALPLLASAGTPEVSILMLDVAGASPTDYVNVVNACLNVQSCVGITVMGVADPDSWRAS
HG259   211 HLSAGQGAGVLQALPLLASAGTPEVSILMLDVAGASPTDYVNVVNACLNVQSCVGITVMGVADPDSWRAS
HG378   211 HLSAGQGAGVLQALPLLASAGTPEVSILMLDVAGASPTDYVNVVNACLNVQSCVGITVFGVADPDSVRAS
HG493   211 HLSAGQGAGVLQALPLLASAGTPEVSILMLDVAGASPTDYVNVVNACLNVQSCVGITVFGVADPDSTRAS
HG603   211 HLSAGQGAGVLQALPLLASAGTPEVSILMLDVAGASPTDYVNVVNACLNVQSCVGITVFGVADPDSVRAS
HG630   211 HLSAGQGAGVLQALPLLASAGTPEVSILMLDVAGASPTDYVNVVNACLNVQSCVGITVMGVADPDSVRAS
HG649   211 HLSAGQGAGVLQALPLLASAGTPEVSILMLDVAGASPTDYVNVVNACLNVQSCVGITVMGVADPDSVRAS

        281 .......290.......300.......310........
HG3     281 TTPLLFDGNFNPKPAYNAIVQDLQQGSIEGRGHHHHHH
HG44    281 TTPLLFDGNFNPKPAYNAIVQDLQQGSIEGRGHHHHHH
HG57    281 TTPLLFDGNFNPKPAYNAIVQDLQQGSIEGRGHHHHHH
HG185   281 TTPLLFDGNFNPKPAYNAIVQDLQQGSIEGRGHHHHHH
HG198   281 TTPLLFDGNFNPKPAYNAIVQDLQQGSIEGRGHHHHHH
HG259   281 TTPLLFDGNFNPKPAYNAIVQDLQQGSIEGRGHHHHHH
HG378   281 TTPLLFDGNFNPKPAYNAIVQDLQQGSIEGRGHHHHHH
HG493   281 TTPLLFDGNFNPKPAYNAIVQDLQQGSIEGRGHHHHHH
HG603   281 TTPLLFDGNFNPKPAYNAIVQDLQQGSIEGRGHHHHHH
HG630   281 TTPLLFDGNFNPKPAYNAIVQDLQQGSIEGRGHHHHHH
HG649   281 TTPLLFDGNFNPKPAYNAIVQDLQQGSIEGRGHHHHHH
```

**Supplementary Figure 7. Multiple sequence alignment of HG-series Kemp eliminases.**

```
                    1 ........10........20........30........40........50........60........70
KE70                1 MTDLKASSLRALKLMHLAT-ANDDDTDEKVIALCHQAKTPVGTTDAIYIYPRFIPIARKTLKEQGTPEIR
KE70-S20a/A240a     1 MTDLKASSLRALKLMHLATSANDDDTDEKVIALCHQAKTPVGTTDAIYIYPRFIPIARKTLKEQGTPEIR
R2_7/12F            1 MTDLKASSLRALKLMHLAT-ANGDDTDEKVIALCHQAKTPVGTTDAIFIYPRFIPIARKTLKEQGTPEIR
R4_8/5A             1 MTDLKASSLRALKLMHLATSANDDDTDENVIALCHQAKTPVGTTDAIFIYPRFIPIARKTLKEQGTPEIR
KE701               1 MTDLKASSLRALKLMDLATSANDDDTDEKVIALCHQAKTPVGTTQAIFIYPRFIPIARKTLKEQGTPEIR
KE703               1 MTDLKASSLRALKLMDLATSANDDDTDEKVIALCHQAKTPVGTTNAIFIYPRFIPIARKTLKEQGTPEIR
KE703b              1 MTDLKASSLRALKLMHLATSANDDDTDEKVIALCHQAKTPVGTTDAIFIYPRFIPIARKTLKEQGTPEIR

                   71 ........80........90.......100.......110.......120.......130.......140
KE70               70 IWTSTNFPHGNDDIDIALAETRAAIAYGADGVAVVFPYRALMAGNEQVGFDLVKACKEACAAANVLLSVI
KE70-S20a/A240a    71 IWTSTNFPHGNDDIDIALAETRAAIAYGADGVAVVFPYRALMAGNEQVGFDLVKACKEACAAANVLLSVI
R2_7/12F           70 IWTSTNFPHGNDDIDIALAETRAAIAYGADGVAVVFPYRALMAGNEQVGFDLVKACKEACAAANVLLSVI
R4_8/5A            71 ICTSTNFPHGNDDIDIALAETRAAIAYGADGVAVVFPYRALMAGNEQVGFDLVKACKEACAAANVLLSVI
KE701              71 ICTSTNFPHGNDDIDIALAETRAAIAYGADSVAVVFPYRALMAGNEQVGFDLVKACKEACAAANVLLQVI
KE703              71 IVTSTNFPHGNDDIDIALAETRAAIAYGADMVAVVFPYRALMAGNEQVGFDLVKACKEACAAANVLLQVI
KE703b             71 IVTSTNFPHGNDDIDIALAETRAAIAYGADMVAVVFPYRALMAGNEQVGFDLVKACKEACAAANVLLQVI

                  141 .......150.......160.......170.......180.......190.......200.......210
KE70              140 IETGELKDEALIRKASEISIKAGADHIVTSTGKVAVGATPESARIMMEVIRDMGVEKTVGFIPAGGVRTA
KE70-S20a/A240a   141 IETGELKDEALIRKASEISIKAGADHIVTSTGKVAVGATPESARIMMEVIRDMGVEKTVGFIPAGGVRTA
R2_7/12F          140 IETGELKDEALIRKASEISIKAGADHIVTSTGKVAVGATPESARIMMEVIRDMGVEKTVGFIPAGGVRTA
R4_8/5A           141 IETGELKDEALIRKASEISIKAGADYIVTSTGKVAVGATPESARIMMEVIRDMGVENTVGFIPVGGVRTA
KE701             141 IETGELKDEALIRKASEISIKAGADNIATSTGKVAVGATPESARIMMEVIRDMGVEKTVGFIPVGGVRTA
KE703             141 IETGELKDEALIRKASEISIKAGADNIATSTGKVAVGATPESARIMMEVIRDMGVEKTVGFIPVGGVRTA
KE703b            141 IETGELKDEALIRKASEISIKAGADNIATSTGKVAVGATPESARIMMEVIRDMGVEKTVGFIPVGGVRTA

                  211 .......220.......230.......240.......250.......260.......270.
KE70              210 EDAQKYLAIADELFGADWADARHYAFGAS-SLLASLLKALGHGDGKSASSYGSLEHHHHHH
KE70-S20a/A240a   211 EDAQKYLAIADELFGADWADARHYAFGASASLLASLLKALGHGDGKSASSYGSLEHHHHHH
R2_7/12F          210 EEAQKYLAIADELFGADWADARHYAFGAS-SLLASLLKALGYGDGKSASSYGSLEHHHHHH
R4_8/5A           211 EDAQKYLAIADELFGADWADARHYAFGASGSLLASLLKALGHGDGKSASSYGSLEHHHHHH
KE701             211 EDAQKYLAIADELFGADWADARHYAFGASASLLASLLKALGHGDGKSASSYGSLEHHHHHH
KE703             211 EDAQKYLAIADELFGADWADARHYAFGASASLLASLLKALGHGDGKSASSYGSLEHHHHHH
KE703b            211 EDAQKYLAIADELFGADWADARHYAFGASASLLASLLKALGHGDGKSASSYGSLEHHHHHH
```

**Supplementary Figure 8. Multiple sequence alignment of KE70-series Kemp eliminases.**
